# The role of pigments in light color variation of the firefly *Photinus pyralis*

**DOI:** 10.1101/2024.09.23.614534

**Authors:** M.S. Popecki, R.L. Rogers, S.A. Archer-Hartmann, J. P. Wares, K.F. Stanger-Hall

## Abstract

Fireflies use bioluminescent signals to communicate with their mates. Luciferase has been thought to be the sole contributor to light color; however, populations of the Photinus pyralis firefly display variation in the color of their emitted signals yet have identical luciferase sequences. Here, we examined whether pigments could be present in the light organs of the twilight-active species P. pyralis and contribute to this variation. We detected patterns of expression that suggest ommochrome and pterin screening pigments are expressed in P. pyralis light organs and could filter light emitted by luciferase and play a role in signal tuning. There were no significant differences between the pigment gene expression of P. pyralis individuals with yellower and greener signals. Our study provides alternative mechanisms that could influence pigments in P. pyralis light organs that could also play a role in modifying signal color.

## Introduction

Fireflies (beetles in the family Lampyridae) are famous for their bioluminescent displays during summer nights when they search for a conspecific mate. There are more than 150 firefly species in North America and 10 or more firefly species may display at a single location (Lloyd, 1969). To recognize conspecifics and reduce mating mistakes, fireflies emit signals in a species-specific pattern that includes flash duration, inter-flash intervals and repeat patterns (e.g., Stanger-Hall & Lloyd, 2015). There is also a diversity in firefly light color across species, ranging from green to orange (550-579 nm; Hall et al., 2016). While light color may not have a direct role in mate choice (Loyd, 1966), it is crucial for signal detection (i.e., generating contrast with the visual background) and thus a prerequisite for mating success. The evolution of firefly light color has been postulated as a two-step process for improving the detection of firefly light signals in their signaling environment (Seliger 1982a,b).

Ancestral fireflies most likely emitted green bioluminescence during their nocturnal displays (Oba et al., 2020), matching the peak green sensitivity of insect eyes (Briscoe & Chittka, 2001). However, when some fireflies (i.e., many species in the genus *Photinus*) shifted their activity time from dark to twilight, possibly due to predation pressure from nocturnal *Photuris* fireflies (Lloyd, 1973), green light signals were more difficult to detect by potential mates among the now visible green vegetation. This likely introduced a selective pressure on firefly eyes, resulting in a shifted peak visual sensitivity from green towards yellow (i.e., to detect the yellow wavelengths within the green light signals), increasing the visual contrast between signal and background environment (Seliger 1982a,b). As the visual sensitivity shifted towards yellow, fireflies that emitted light spectra with relatively more yellow wavelengths had increased mating success as they were more easily seen by potential mates (Figure 1). Such shifts in peak visual sensitivity towards more yellow light, followed by a corresponding shift in signal color in different firefly species, diversified the range of light color in twilight-active fireflies from green-yellow to orange (Lall et al., 1980; Seliger 1982a,b; Hall et al., 2016).

**Figure 1.**
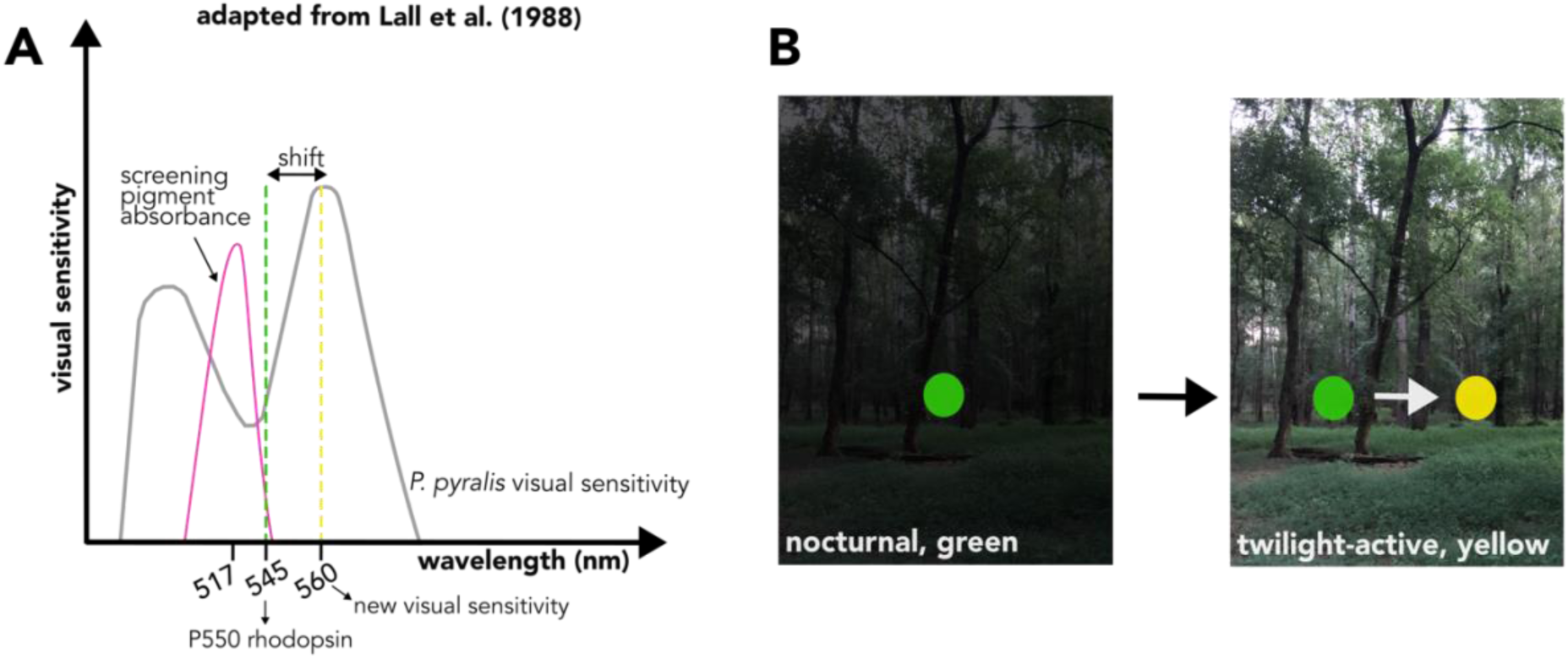
Diagram of the shifted peak visual sensitivity in *P. pyralis*. An unknown screening pigment absorbs wavelengths at the green end of the incoming light spectrum (Lall et al., 1988; Cronin et al., 2000), resulting in relatively more yellow light stimulating the P550 rhodopsin (with LW opsins: Sander & Hall 2015), which maximally absorbs photons at 545 nm (green). As a result, the peak visual sensitivity has narrowed from a broad green/yellow to a narrow, yellow-shifted peak sensitivity (modified from Lall et al., 1988). **(A)** Within the photic cells of the light organ, bioluminescence is produced by luciferase. As the light passes through pigment granules in the photic cells, wavelengths at the green end of the spectrum are absorbed, which shifts the emitted light color towards yellow. **(B)** At twilight, yellow or orange signals are more conspicuous than green signals due to their higher contrast with visible background foliage.

Our study focuses on the second step, the evolution of firefly light color. Adult fireflies generate light signals within their light organs, which are specialized abdominal segments with a clear cuticle. While many components of the bioluminescent reaction, such as the luciferase enzyme substrate luciferin, are conserved among beetles (Seliger et al., 1961; Seliger & McElroy, 1964), firefly luciferase exhibits structural diversity (Hastings, 1983). The amino acid sequence of luciferase determines enzymatic activity, and thus bioluminescent properties such as light intensity (Liu & Urban, 2017) and color (Nakatsu et al., 2006). *In vitro* studies of luciferase have demonstrated that individual amino acid substitutions can alter these properties (Fujii et al., 2007; Kajiyama & Nakano, 1991; Branchini et al., 1999; Branchini et al., 2017). Under the traditional paradigm, luciferase is thought to be the sole determinant of bioluminescence color in fireflies, however this was challenged by Lower et al. (2018), who documented *Photinus pyralis* populations with differences in emitted light color but identical luciferase amino acid sequences.

This raises the question: how is variation in firefly light color generated? Here, we propose and test the hypothesis that pigments in the light organ are contributing to the emitted light color in fireflies. We predict that (1) pigments are present in the light organs of fireflies, i.e., genes encoding enzymes and transporters in pigment biosynthesis pathways are expressed, and that (2) these pigments can modify firefly light color by absorbing part of the luciferase light spectrum and reflecting other wavelengths, thus effectively shifting the light color that is emitted from the light organ. Indirect evidence for light organ pigments comes from observations of light organ hue changes immediately before onset of firefly activity. To the human eye, the light organs of inactive *Photuris* and *Photinus* fireflies appear ivory to pale-yellow in color. However, right before they become active at twilight, the light organs of *Photinus* species such as *P. australis* and *P. pyralis* undergo a light organ hue change to a deeper yellow or yellow-orange (Figure 2). After their light displays end each evening and fireflies return to rest, their light organs revert to their paler color. Twilight active *P. scintillans* undergo a similar change with a hue change from yellow to pink (Faust et al., 2019).

**Figure 2.**
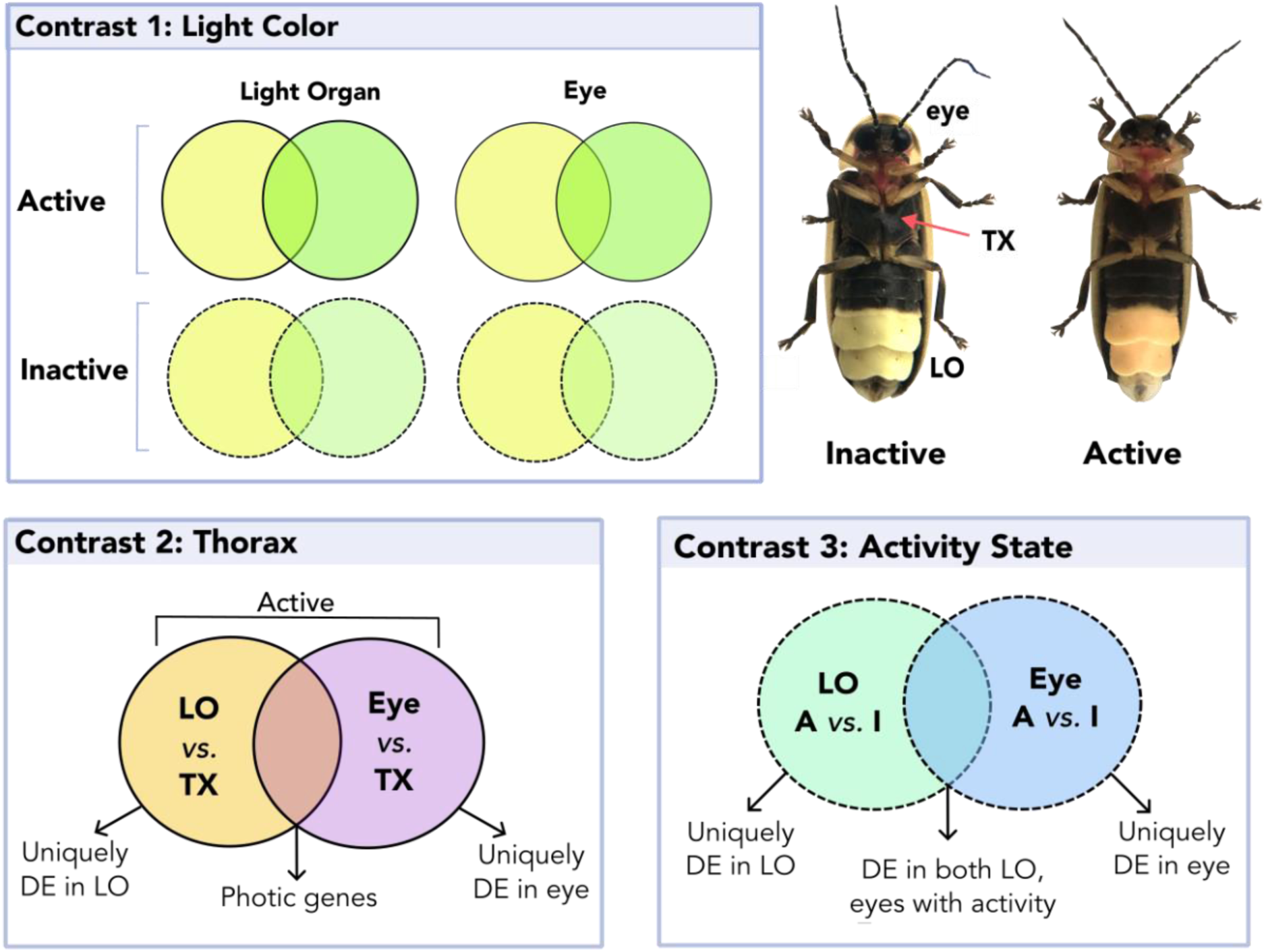
Overview of differential gene expression analyses. LO=Light organ, TX=Thorax. We compared expression patterns between tissues, activity state, and light color. Note: The light organ of *Photinus pyralis* fireflies appears ivory-yellow during the inactive state (left) and shifts to a yellow-orange hue while fireflies are active (flying and signaling) at twilight (right).

This rapid and reversible color change of the light organ suggests the presence of pigments in the light organs, which could either be newly generated and/or transported into pigment granules below the clear cuticle of the light organ before fireflies become active. Morphological studies of the firefly light organ have documented “differentiated-zone granules” within the photophore, located at the photic cell borders in proximity to mitochondria and peroxisomes (Peterson & Buck, 1968, Ghiradella & Schmidt, 2004). This description is consistent with the presence of pigments, which are synthesized in granules known as lysosomal-related organelles (LROs). Pigment biosynthesis pathways consist of a series of enzymes that produce pigment precursors. The precursors produced in the later steps of the pathways are transported into granules, where they can undergo additional modifications that result in mature pigments with specific light absorption characteristics (Figon & Casas, 2019; Andrade & Carneiro, 2021). Wavelengths that are not absorbed are reflected, giving different pigments their different colors.

To test our hypothesis that pigments in the light organ of *Photinus pyralis* contribute to the variation in light color within and between *P. pyralis* populations with identical luciferases, we first determined whether pigment genes and/or pigment transporter genes are expressed in *P. pyralis* light organs. Color generation by pigments has been described for several pigment classes, the most common of which include melanins, carotenoids, pterins and ommochromes (Andrade & Carneiro, 2021). Unlike melanins, which are frequently used for cryptic coloration across animals, and carotenoids, which rely on a dietary uptake, pterins and ommochromes are produced endogenously (Andrade & Carneiro, 2021). Due to their universal role in insect pigmentation, our emphasis will be on pigments in the pterin and ommochrome pathways. Pterins (appearing red or yellow to human eyes) and ommochromes (appearing yellow, red, brown, or violet) are common pigments in insects and protect against oxidative damage induced by UV exposure and metabolic toxins and are also integral for visual acuity in insect eyes (Insausti et al., 2013). Since pigments are stored in granules, we will also examine whether genes involved in granule formation and/or transport of precursors into granules are expressed.

To determine if pigments influence light color, we tested if these pigment, granule, and transporter genes differ in type or expression level between the light color extremes of *P. pyralis* (greener light: 560-562 nm versus yellower light: 565-568 nm). Since pigment expression and/or transport may change prior to signaling activity (as indicated by the hue change of light organs), we will compare pigment gene expression in the light organ between active (dusk) and inactive (morning) fireflies to narrow down gene candidates likely involved in shifting light organ hue and possibly light color. On the other hand, pigments may be generated continuously, and transport processes may be responsible for the observed hue change, therefore we will also identify any pigment genes expressed in light organs, even if their expression does not change with activity. Since screening pigments are known to play an important role in insect eyes (Ziegler, 1961; Linzen, 1967; Langer, 1975), including fireflies (Lall et al. 1988, Cronin et al., 2000), we will examine whether any expressed pigment genes and/or transporters in firefly light organs are also expressed in firefly eyes, thus providing a direct pigment-based mechanism for the reported tuning between emitted light color (light organ) and the visual sensitivity of firefly eyes (Lall et al., 1980, Seliger 1982a,b). In addition, we examine whether the bright red/yellow aposematic coloration of the firefly headshield (with unknown pigments) shares any pigments with firefly light organs and/or eyes.

## Methods

### Field collections

We collected *Photinus pyralis* fireflies from two natural populations in Georgia with light color variation in an urban area of Athens, GA (-83.369901, 33.96541; greener light color, Flint Street) and a farm located in Watkinsville, GA (-83.4009, 33.8214; yellower light color, Rose Creek) located about 20 km (straight line distance) from each other (Figure S1). After capture with a net or by hand, we immediately recorded the light spectra of firefly flashes using a portable Jaz spectrophotometer (Ocean Optics Jaz, Orlando, FL) with a laptop computer (SpectraSuite 2.0). Fireflies were either immediately flash-frozen on dry ice for subsequent preservation in RNAlater-ICE (for gene expression of active fireflies) or kept overnight at room temperature in Falcon tubes (with small pieces of apple and damp coffee filters to prevent dehydration) before flash-freezing them 12 hours after their activity period (for gene expression of inactive fireflies). All specimens were stored in a -80°C freezer until transitioning into RNAlater-ICE following manufacturer protocol (Thermofisher, Waltham, MA) with storage at -30°C until RNA isolation.

### Measuring firefly light color

Firefly light spectra collected in the field were analyzed using a custom Mathematica script by fitting a cubic polynomial to raw measurements (Hall et al., 2016). Measurements with a saturated amplitude (light intensity) were removed, as their peak emission wavelengths could not be determined accurately. For each firefly, we calculated the mean wavelength (at peak light intensity) across its individual recorded light spectra, as well as the standard deviation and range (Table S1), using a custom script (Hall et al. 2016). For transcriptome analysis, we chose the eight fireflies (biological replicates) from each population with the most extreme light colors: 8 with the most yellow (566-569 nm) flashes from our yellower population and 8 with most green (560-562 nm) flashes from our greener *P. pyralis* population. Four of the eight fireflies in each color sample were preserved in their active state and four in their inactive state. For pigment analysis, we used three replicates for each state (Table S2; Supplement).

### RNA isolation and sequencing

For each preserved specimen, we dissected the eyes (light sensing), light organs (light generation) and thorax muscle (reference tissue not associated with light signaling). Light organs included the clear ventral cuticle, the layer of photic cells where bioluminescence is produced, and the reflective layer. To generate a reference transcriptome for a tissue with conspicuous pink body pigmentation, we extracted RNA from the head shield (pronotum). As this tissue was sparse and fatty, pooling of N=3 active individuals achieved an RNA concentration high enough for sequencing. RNA was isolated from all tissues using Trizol protocol (https://github.com/mspopecki/Gene-expression-Photinus-pyralis.git). All samples were subjected to DNAse treatment (TURBO Invitrogen DNAse) and potential residual organic contamination was removed (Zymo Clean & Concentrator-25) before a quality check (Qubit and Bioanalyzer) and transcriptome sequencing (150-PE, non-strand specific) at Novogene (Sacramento, CA). The head shield sample was sequenced on a separate run (Table S3).

### Bioinformatic processing from reads to gene abundance

Transcript reads were assessed for quality using FASTQC (0.11.9) with minor trimming using default parameters of TrimGalore! (0.6.5). Trimmed reads were aligned to the *P. pyralis* genome (version 1.4, Fallon et al. 2018) with STAR (2.7.3a) and expression abundance was calculated with RSEM (1.3.3). To ensure libraries had high quality and that missing data would not reduce our power to evaluate changes in gene expression, we removed any transcriptomes with fewer than 10% the average library size from subsequent analyses. This resulted in the removal of one transcriptome (FS34I3: inactive light organ, greener light color) (Table S3).

### Experimental Design

Our approach to contrasting expression patterns across samples and tissues is illustrated in Figure 2. First, we use light color of the individual (categorized as yellow or green) to contrast the activity state within photic tissues (Contrast 1). Second, we contrast signaling tissues against non-signaling (thorax) tissues pooled from all samples to determine genes with significantly different activity from the thorax (Contrast 2), as those genes that are shared between LO and eyes in being distinct from thorax may have a more important role in light signaling. Finally, we reevaluated activity state using pooled tissues in light organs and eyes with a focus on differential expression of recognized pigment genes (Contrast 3).

### Analysis of differential gene expression

We identified variation in gene expression between tissues and/or activity states using the R package DESeq2 (1.36.0). We filtered genes with low or no expression by requiring at least three samples to have an expression level of ≥3 transcripts per million (TPM) for that gene. Significance values were adjusted with the Benjamini-Hochberg correction for false discovery rate (Benjamini & Hochberg, 1995) in multiple comparisons. We filtered differentially expressed pigment genes from transcripts with an adjusted p-value < 0.05. To investigate patterns of differential expression across all transcripts, for contrasts with thorax (Contrast 2) we focused on those with both an adjusted p-value < 0.05 and log_2_ fold change greater than |2| but relaxed the log_2_ fold change threshold to |1| when comparing light color (Contrast 1) and activity state (Contrast 3) to investigate subtler changes within tissues. Given the threshold for biological relevance of pigment gene expression is unknown in fireflies, we reported all pigment genes that were differentially expressed at p-adjusted <0.05, both with and without the log2 fold change threshold for each contrast.

### Highly expressed genes in the head shield

Since it is currently unknown which pigments are used to produce the pink aposematic coloration on the head shield of fireflies and whether these same pink pigments could be utilized to shift the emitted light color in the light organ, we included the head shield into our pigment expression analyses. We identified the most highly expressed (HE) genes in the firefly head shield for comparison with genes expressed in active light organs by calculating the 90th percentile of expressed transcripts (>3 TPM).

### Identification of firefly orthologs and pigment genes in the *P. pyralis* genome

We used OrthoFinder (v. 2.3.8) with the protein genomes of 13 insect species (Table S4) which obtains all gene families (orthogroups) to identify homologous sequences, including putatively duplicated genes. To identify orthologous pigment genes between model organism *D. melanogaster* and *P. pyralis*, we searched orthogroups for the *D. melanogaster* FlyBase identifier (Table S5). We focused on genes involved in ommochrome and pterin biosynthesis (Figure 3) given their widespread use as screening pigments. To identify the MFS pigment transporter gene for *re* in *P. pyralis,* we used the published AA sequence for *Tribolium castaneum* as query against the *P. pyralis* genome in BLASTP (2.9.0). We chose the best hit to *P. pyralis* (lowest e-value and highest bit score) to identify the most likely ortholog of this gene in *P. pyralis*.

**Figure 3.**
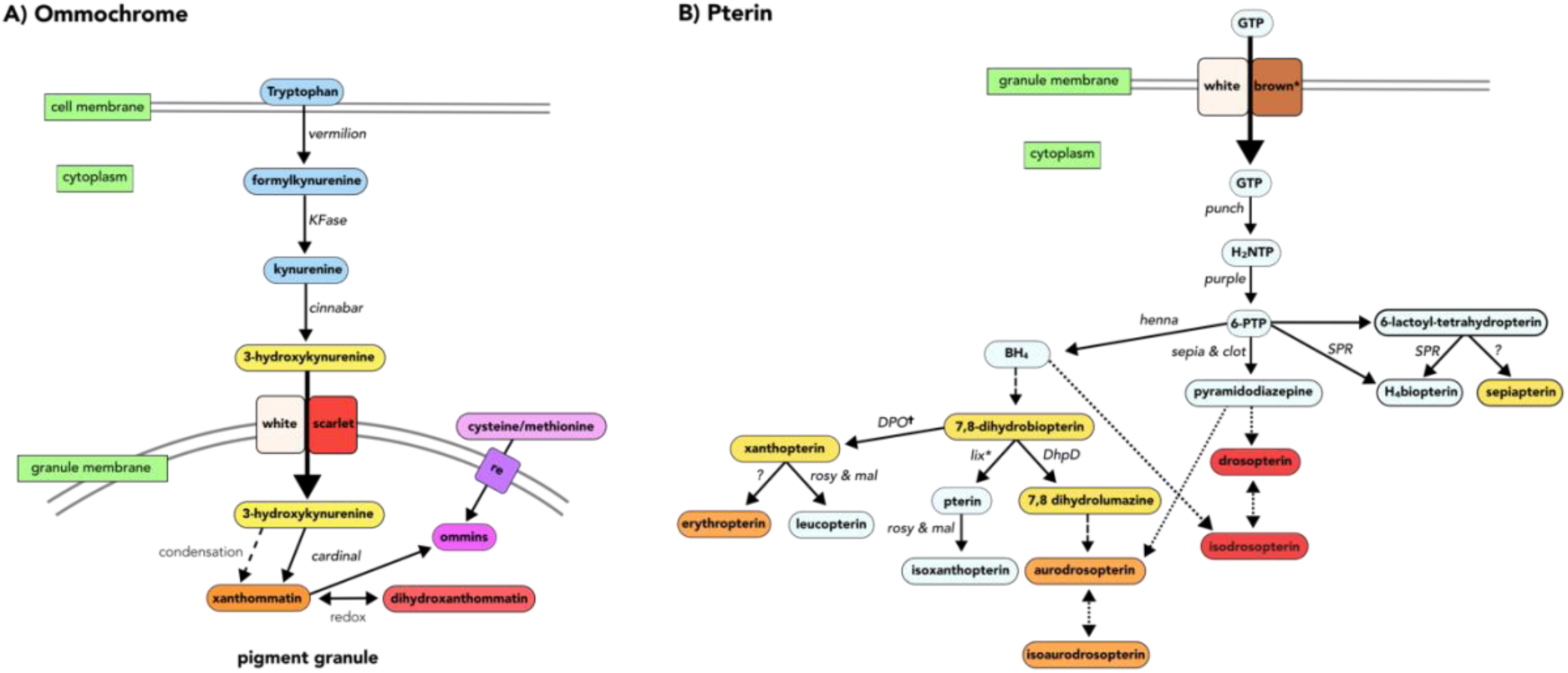
Pigment biosynthesis pathways for (A) ommochromes (modified from Figon & Casas, 2018; Vargas Lowman et al., 2019) and (B) pterins (modified from Vargas Lowman et al., 2019; Andrade & Carneiro, 2021; Ferré, 2024) in *Drosophila melanogaster*. Includes enzymatic (straight lines) and non-enzymatic/condensation (dashed lines) reactions. Boxes: pigment substrates (with pigment color); gene name of the enzyme catalyzing the respective step (arrow) in *D. melanogaster*. *White, scarlet*, *brown*: ABC transporters involved in ommochrome and pterin synthesis; *red egg (re)*: MFS transporter in ommochrome pathway; (*)genes we did not identify (< 3TPM) in *P. pyralis* (Table S5).

### Weighted co-expression gene network analysis (WCGNA) for light organs

To identify genes with shared expression patterns, we complemented our differential expression analyses with a weighted co-expression gene network analysis using 15 light organ samples (4 active yellow, 4 active green, 4 inactive yellow, 3 inactive green; Figure S2) (R package WCGNA v. 1.72-1). We prepared the transcript count data for this network analysis with the “getVarianceStabilizedData” function from DESeq2, a homoscedastic transformation used to normalize variance across samples for clustering (Langfelder & Horvath, 2008).

We generated a signed network, which considers only positive correlations between light organ genes (genes that share the same direction of expression such as up or down-regulation, as opposed to repression, where genes are expressed in opposite directions). We used hierarchical clustering to group genes with similar expression patterns (low dissimilarity) into sets of co-expressed genes (modules).

To summarize gene expression within modules, we performed a principal component analysis on module gene expression using the “moduleEigengenes” function. The first principal component, (module eigenvector or ME), was used to describe the expression trends across the genes in the module. To determine if module gene expression (ME) was significantly correlated with the mean recorded light color of the fireflies in each group of samples, we performed a Pearson’s correlation using the “cor” function in base R (4.2.1). To model the correlations between ME with activity state (active, inactive) as a discrete variable, we used a linear model using the “lmFit” function from the limma package (3.52.4) in R. To address the variability in expression across samples (from smaller sample size), we used the limma “eBayes” function to stabilize variance (shift genes with both high and low variation toward the mean variance). The statistics were generated by the limma “topTable” function, which was also used to apply the Benjamini-Hochberg (BH) false discovery rate method correction to p-values to account for multiple comparisons. We report modules with correlations between ME and either light color or activity state with a p-value < 0.05 and adjusted p-value < 0.05 as robust; modules with a p-value <0.05 and an adjusted p-value slightly > 0.05 are reported as “of interest”. Given the limitations of sample size and high variation within natural populations, we included all modules with p<0.05 in our analysis.

### Gene ontology enrichment and module characterization

To provide biological context to significant gene expression modules, we conducted gene ontology (GO) enrichment analyses. To increase the annotation accuracy, we did not solely rely on homology with *D. melanogaster*, but also retrieved *P. pyralis* GO terms using eggNOG-mapper v2 (2.1.9), which compares protein orthology across several databases. We subsequently tested for GO term enrichment using the R package TopGo (2.48.0) using the default algorithm (weight-01) for Fisher’s Exact Test. We used a background of all annotated genes for differential expression across tissues, and genes expressed in LO for module gene lists. Resulting p-values were corrected for multiple testing (BH) with “p.adjust” function from base R stats function (4.2.1). GO terms with p < 0.005 were considered significant. We searched all modules for the presence of pigment genes. As gene annotations related to module function and firefly signal production are sparse, we further characterized our modules by comparing the genes in the different modules to the gene list published by Fallon et al. (2018), referred to as “LO genes.” Modules were tested for enrichment with pigment and bioluminescence genes, in addition to select differentially expressed gene lists using Fisher Exact Test in R; p-values were adjusted for multiple tests with False Discovery Rate (FDR).

### Gene connectivity within modules

To understand the connection of genes within significant modules, we identified highly connected genes (hubs). Hubs are strongly associated with the expression of other genes in the same module and could potentially regulate their gene expression (Yu et al., 2017). We first calculated Module Membership (MM: correlation between expression of a gene and a module eigenvector) using the “signedKME” function. Next, we determined Gene Significance (GS) as the correlation between the expression of an individual gene and the phenotype of interest (i.e., light color, activity state) using Pearson’s correlations. Following Langfelder & Horvath (2008), genes with MM > 0.8 and absolute value of GS > 0.2 with p-adjusted (BH) <0.05 were identified as hubs. To test the relationship between gene expression and activity state, we determined GS using a linear model (limma) and established hubs using the same thresholds.

### Pterin analysis of signaling tissues

To supplement the data from our gene expression analysis with a pigment analysis in light organs, eyes, and HS, we extracted pigments and used UV-vis followed by LC-MS in a preliminary pigment analysis to identify the most abundant pigments present in the LO and eye extracts of *P. pyralis* fireflies and to inform future more-in depth studies (Supplement).

## Results

### Field collections

We collected *P. pyralis* from two populations (20 km apart) with a difference in mean light color. To rule out evolutionary divergence as a factor, we also sequenced a mitochondrial COI gene region (∼620 bp) (Figure S3) for all individuals and estimated Hudson’s Snn statistic for differentiation of subpopulations (Hudson, 2000); sequences are deposited at NCBI (accession numbers SAMN38520475-38520511). Mitochondrial (Snn=0.5; p ∼1) and additional whole-genome resequencing data confirmed no measurable evolutionary divergence of these populations (Popecki, Rogers, et al., *in prep*). The uniformity of luciferase amino acid sequences was confirmed by alignment (Figure S4).

### Firefly light color

We calculated the mean light color for each firefly as mean wavelength (at highest light intensity) across all their recorded light spectra (Table S1, Figure 4), which were used to determine population means (grouped by activity). The mean (x̄ ± std. error) emitted light color of the four fireflies preserved in the active state was 566.89±0.51 nm for the yellow emitting fireflies and 560.95±0.19 nm for the green emitting fireflies. The mean emitted light color of the fireflies preserved in the inactive state was 566.17±0.74 nm for the four yellow emitting fireflies and (after removing the sample with a small library size: FS34I3) and the mean was 560.82±0.16 nm for the three remaining green fireflies.

**Figure 4.**
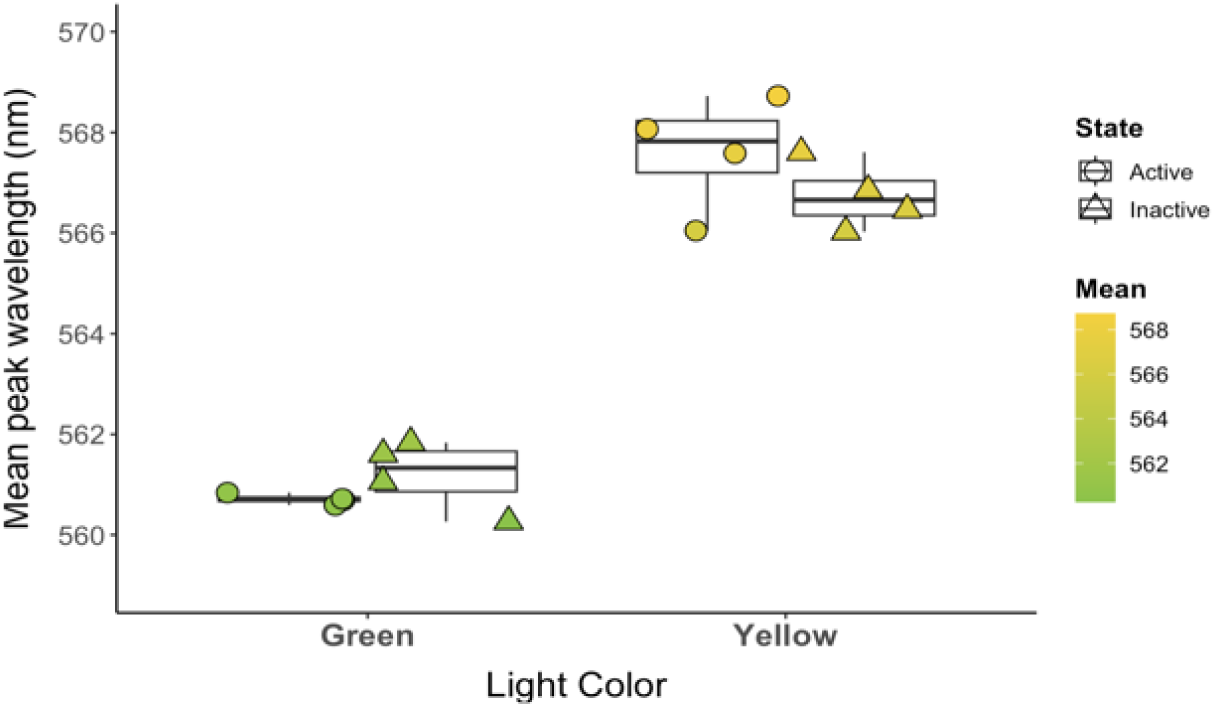
Mean light color (wavelength at peak intensity in nm: x̄ ± std. error) of fireflies from green and yellow populations chosen for transcriptome analysis. Mean: wavelength (nm) as it corresponds to light color perceived by the human eye. State: fireflies used for gene expression analysis and preserved in active (circle) or inactive (triangle) state.

### Transcriptomes

We sequenced multiple tissues from a total of 16 fireflies for transcriptomic analysis (4 active and 4 inactive for each light color category) to generate a total of 38 transcriptomes across light colors, activity states and tissues (Table S3) with a mean library size (x̄± std. dev.) of 10,128,638 ±1,665,770 read counts. Small library size due to missing data can bias results, so we removed samples with less than 10% of mean library size resulting in the exclusion of one inactive light organ replicate (green: FS34I3 with 717,767 reads). Sequences are archived at the NCBI Short Read Archive (accession 38520475-38520511). Eliminating genes with low expression (< 3 TPM) in at least three samples per tissue group resulted in 12,590 (from 15,764) transcripts in our final analyses.

### Pigment, granule, and transporter genes in the genome of *P. pyralis*

To identify firefly orthologs of pigment-related genes in *D. melanogaster* and *T. castaneum*, we searched orthogroups (gene families) for 39 pigment genes, including pterins (with their purine precursors), ommochromes, transporter and granule genes (Table S5). Overall, 53 pigment genes from 34 orthogroups (includes multiple copies) were identified in *P. pyralis* (Table 1, Figure 5; Table S6), though six pigment genes (*purple1, purple4, white3, white4, cinnabar1, and sepia4*), were not expressed (>3 TPM) in any firefly tissue (active and inactive replicates pooled). *Purple3* (pterin) was expressed only in HS, and five pigment genes were absent from the thorax (pterins: *DhpD2, DhpD3, sepia1* and ABC transporter: *scarlet*). Seven orthogroups contained increased copy number (Figure S5-6), with the highest number found within pterins. We did not recover any orthologs for or *lix* (pterin) or *brown* (ABC transporter in pterin pathway).

**Figure 5.**
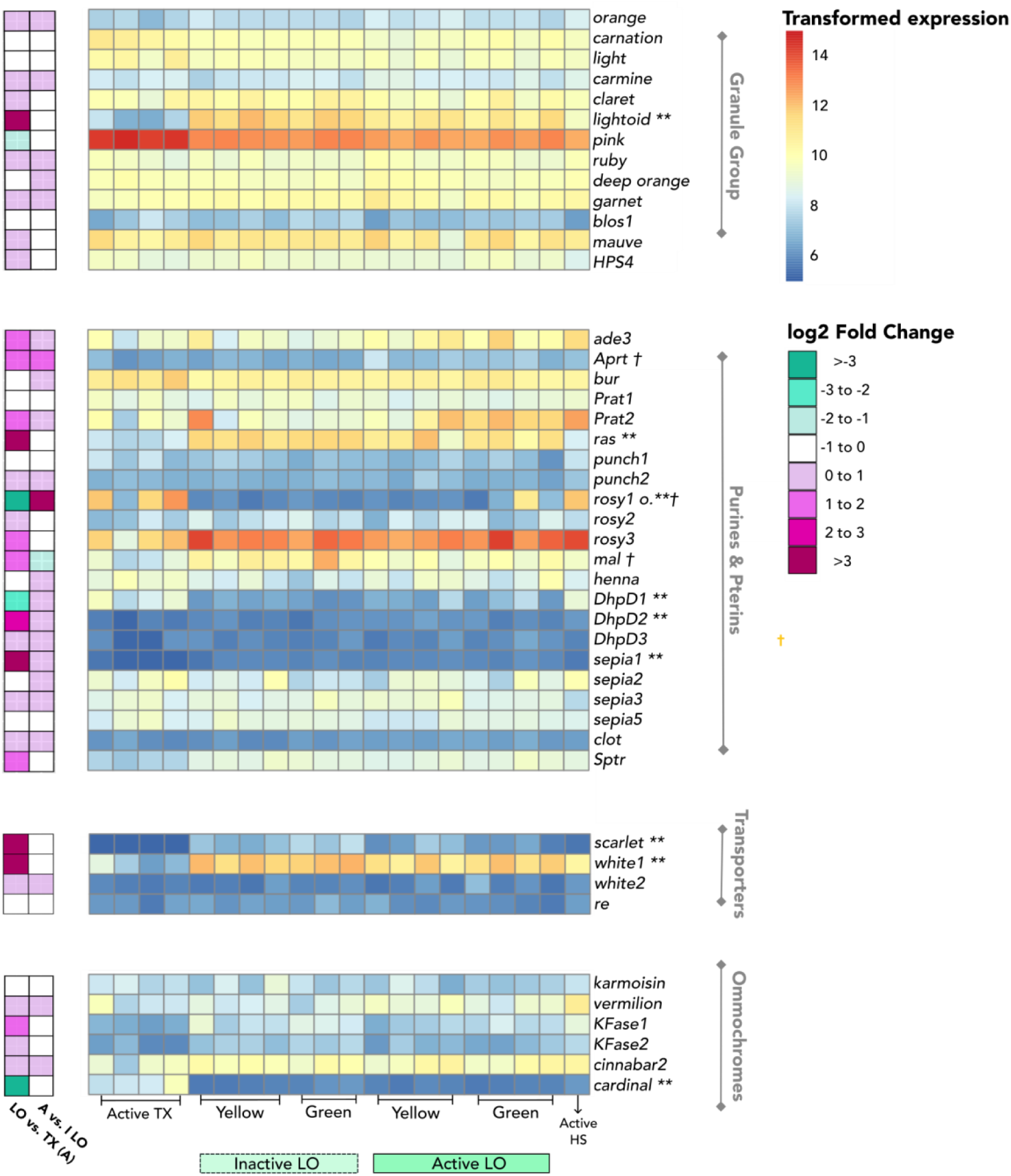
Heatmap (pigment gene expression levels) across samples from active thorax (TX), inactive and active light organs (light organ), and active head shield (HS). For ommochromes, pterins (purines included), the pigment genes are ordered according to their approximate placement in the *D. melanogaster* pigment pathways; transporters are listed in the ommochrome block. The granule group is unordered. Color legend is scaled by TPM with color indicating genes with low (blue), moderate (yellow), or high (red) expression. Pigment genes that were recovered but not expressed > 3 TPM in at least one tissue were omitted. Log2 fold change values are included to visualize magnitude of expression changes in pigment genes. Differential expression significance denoted by ** >|2| log2 fold change (LO vs. thorax) and † > |1| log2 fold change (active vs. inactive LO).

**Table 1:** Pigment genes used to identify orthologs from the *P. pyralis* genome (PPYR ID) (based on list from Croucher et al., 2013). FlyBase ID=Identity of *D. melanogaster* genes and protein ID of *T. castaneum red egg (re)*: unique to beetles.

| Gene Name | Class | PPYR ID | Orthogroup | FlyBase ID |
| --- | --- | --- | --- | --- |
| *scarlet* | ABC transporter | PPYR_04279 | OG0000728 | FBgn0003515 |
| *white1* | ABC transporter | PPYR_04280 | OG0000728 | FBgn0003996 |
| *white2* | ABC transporter | PPYR_04281 | OG0000728 | FBgn0003996 |
| *white3*<sup>†</sup> | ABC transporter | PPYR_04282 | OG0000728 | FBgn0003996 |
| *white4*<sup>†</sup> | ABC transporter | PPYR_14447 | OG0000728 | FBgn0003996 |
| *re*<sup>*</sup> | MFS transporter | PPYR_05423 | OG0005173 | TC013631<sup>1</sup> |
| *blos1* | granule | PPYR_11283 | OG0008505 | FBgn0050077 |
| *carmine* | granule | PPYR_03448 | OG0005216 | FBgn0000330 |
| *carnation* | granule | PPYR_02799 | OG0005324 | FBgn0000257 |
| *claret* | granule | PPYR_04283 | OG0002347 | FBgn0000247 |
| *deep-orange* | granule | PPYR_10566 | OG0004412 | FBgn0000482 |
| *garnet* | granule | PPYR_11202 | OG0004785 | FBgn0030089 |
| *HPS4* | granule | PPYR_14639 | OG0006916 | FBgn0034261 |
| *light* | granule | PPYR_02943 | OG0005849 | FBgn0002566 |
| *lightoid* | granule | PPYR_05587 | OG0006327 | FBgn0002567 |
| *mauve* | granule | PPYR_12819 | OG0002750 | FBgn0043362 |
| *orange* | granule | PPYR_02585 | OG0006036 | FBgn0003008 |
| *pink* | granule | PPYR_09054 | OG0009008 | FBgn0029891 |
| *ruby* | granule | PPYR_09882 | OG0003078 | FBgn0003210 |
| *cardinal* | ommochrome | PPYR_05801 | OG0003614 | FBgn0263986 |
| *cinnabar1*<sup>†</sup> | ommochrome | PPYR_12971 | OG0002355 | FBgn0000337 |
| *cinnabar2* | ommochrome | PPYR_06324 | OG0002355 | FBgn0000337 |
| *karmoisin* | ommochrome | PPYR_05108 | OG0007399 | FBgn0001296 |
| *Kfase1* | ommochrome | PPYR_14911 | OG0001298 | FBgn0031821 |
| *Kfase2* | ommochrome | PPYR_04835 | OG0001298 | FBgn0031821 |
| *vermillion* | ommochrome | PPYR_07427 | OG0002319 | FBgn0003965 |
| *clot* | pterin | PPYR_11797 | OG0006442 | FBgn0000318 |
| *DhpD1* | pterin | PPYR_00483 | OG0001971 | FBgn0261436 |
| *DhpD2* | pterin | PPYR_01709 | OG0001971 | FBgn0261436 |
| *DhpD3* | pterin | PPYR_01710 | OG0001971 | FBgn0261436 |
| *henna* | pterin | PPYR_10049 | OG0004429 | FBgn0001208 |
| *mal* | pterin | PPYR_07656 | OG0004893 | FBgn0002641 |
| *punch1* | pterin | PPYR_14140 | OG0003359 | FBgn0003162 |
| *punch2* | pterin | PPYR_14143 | OG0003359 | FBgn0003162 |
| *purple1*<sup>†</sup> | pterin | PPYR_04028 | OG0002731 | FBgn0003141 |
| *purple2* | pterin | PPYR_09660 | OG0002731 | FBgn0003141 |
| *purple3* | pterin | PPYR_09659 | OG0002731 | FBgn0003141 |
| *purple4*<sup>†</sup> | pterin | PPYR_15506 | OG0002731 | FBgn0003141 |
| *rosy1* | pterin | PPYR_11236 | OG0000709 | FBgn0003308 |
| *rosy2* | pterin | PPYR_11237 | OG0000709 | FBgn0003308 |
| *rosy3* | pterin | PPYR_11235 | OG0000709 | FBgn0003308 |
| *sepia1* | pterin | PPYR_09852 | OG0000366 | FBgn0086348 |
| *sepia2* | pterin | PPYR_13566 | OG0000366 | FBgn0086348 |
| *sepia3* | pterin | PPYR_02423 | OG0000366 | FBgn0086348 |
| *sepia4*<sup>†</sup> | pterin | PPYR_14507 | OG0000366 | FBgn0086348 |
| *sepia5* | pterin | PPYR_14121 | OG0000366 | FBgn0086348 |
| *Sptr* | pterin | PPYR_12709 | OG0008551 | FBgn0014032 |
| *ade3* | purine | PPYR_13352 | OG0002091 | FBgn0000053 |
| *Aprt* | purine | PPYR_00822 | OG0006160 | FBgn0000109 |
| *bur* | purine | PPYR_12914 | OG0005692 | FBgn0000239 |
| *Prat1* | purine | PPYR_03533 | OG0001300 | FBgn0004901 |
| *Prat2* | purine | PPYR_10414 | OG0001300 | FBgn0004901 |
| *ras* | purine | PPYR_05313 | OG0003143 | FBgn0003204 |

<sup>†</sup>Was not expressed in any tissue (LO, eyes, thorax) > 3 TPM.
<sup>1</sup>Gene derived from *Tribolium castaneum* (Osanai-Futahashi et al., 2012).

### Pigment genes expressed in light organs (LO) and eyes *of P. pyralis* fireflies

Of the 53 pigment genes in the pterin and ommochrome pathways identified in the *P. pyralis* genome, 46 were expressed (>3TPM) in the LO of *P. pyralis* fireflies (Table S6) and thus could potentially contribute to the formation of colored pigment products in the LO that filter and shift light color. In *P. pyralis* eyes, 46 pigment genes were expressed (>3TPM) (Table S6) and may have a role in tuning of visual sensitivity of firefly eyes. To identify the most likely candidates for shifting light color in LO and for tuning between light spectra and visual sensitivity, we used three contrasts to identify pigment genes that were differentially expressed between light colors, tissue type and activity. To identify the pigment genes with the most substantial differential expression, we used a log2fold change (>|2| for thorax, >|1| with activity and light color within LO and eyes), but since any pigment present may impact light color and tuning, we also report log2fold changes below these thresholds.

### Contrast 1: Light Color

To test our hypothesis that pigments in the LO contribute to the emitted light color in fireflies, we examined differences in pigment gene expression between the LO of *P. pyralis* with yellow and green signals. There were 88 differentially expressed genes (21 upregulated in yellow, 67 upregulated in green), while the same contrast with inactive LO yielded 19 differentially expressed genes (14 upregulated in yellow, 5 upregulated in green). Only *rosy1* (outlier) exhibited differential expression between yellow and green LO (more highly expressed in active green LO; Figure S8).

To identify parallel changes in eyes, we compared pigment gene expression in the eyes of fireflies with “yellow” or “green” flashes. During activity, 105 genes were differentially expressed (77 upregulated in yellow, 28 upregulated in green) between the eyes of green and yellow fireflies. One pigment gene, *sepia3*, was upregulated in the eyes of active fireflies with yellow light (log2fold change=1.022). In inactive eyes, only five genes were differentially expressed (all 5 upregulated in yellow versus green); none of these were pigment genes.

### Contrast 2: Tissues (active state)

Given that tissue type was the primary driver of gene expression differences across samples (Figure S7), and our expectation that pigment genes most relevant to light signals should be more highly expressed in photic tissues (light organs and eyes) compared to non-signaling tissues (thorax), we contrasted the LO and eyes with the thorax of active fireflies. We report all significantly differentially expressed pigment genes (and whether they differed at a log2fold change of >2 or <2).

### Active LO vs. thorax

There were 2,037 differentially expressed genes between active light organs and thorax (1,260 upregulated in light organ, 777 upregulated in thorax) (Figure 6A). Of the 1,260 genes more highly expressed in light organs, six were pigment genes, including 2 pterins (*DhpD2*, *sepia1)*, 2 transporters (*scarlet, white1*), and 1 granule gene (*lightoid*) (Table S7). Relative to thorax, three pigment genes were downregulated in LO (2 pterins: *DhpD1, rosy1*, and 1 ommochrome: *cardinal)*; however, the expression of *rosy1* was driven by one sample (outlier) (Figure S8). In addition, there were 10 pigment genes with significant differential expression (log2fold change < |2|) that were upregulated in active light organs (and four downregulated) (Table S7) and could affect light color. These included 3 pterins (*mal*, *rosy3, Sptr*) and 3 ommochromes (*cinnabar2, KFase1, KFase2*). Significant KEGG pathways are reported in Table S8.

**Figure 6:**
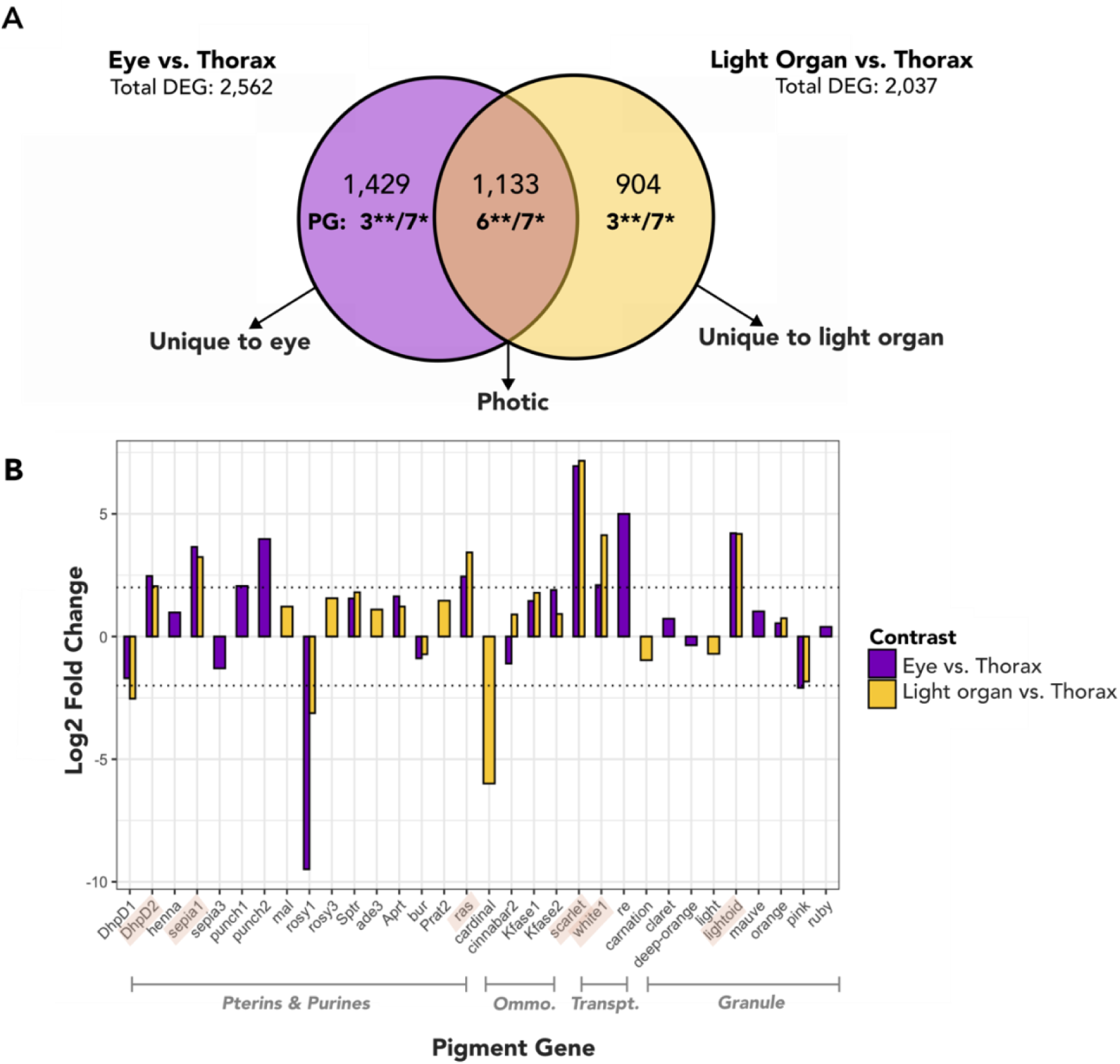
Pigment genes expressed in both light organs and eyes that differ from thorax. **A:** Results of Contrast 1: Comparison of differential gene expression between tissues (active state). **Significant p-adjusted (BH) and log2fold change > |2|, *Significant p-adjusted (BH) DEG. **B:** Expression patterns of 28 genes in the light organs and eyes of active fireflies that were differentially expressed from thorax. Apart from *cinnabar2*, all other pigment genes DE with thorax in both LO and eyes show the same directionality of expression. Dotted lines on y-axis (at 2 and -2 log2 fold change) indicate significance thresholds for Contrast 2: (1) log2 fold change > |2| and p-adjusted < 0.05, (2) p-adjusted < 0.05. Highlighted pigment gene names on x axis indicate **Significant p-adjusted (BH) and log2fold change > |2|.

### Active eye vs. thorax

Between active eyes and thorax, there were 2,562 differentially expressed genes (in eyes: 1,503 upregulated, 1,059 downregulated). Of these, nine pigment genes were upregulated in eye (4 pterins: *DhpD2, sepia1, punch1, punch2*, 2 ABC transporter: *scarlet, white1*, 1 MFS transporter: *re*, and 2 granule genes: *lightoid, ras*), whereas two were downregulated in active eyes: 1 pterin (*rosy1*), and 1 granule gene (*pink*). In addition, there were 14 pigment genes (9 upregulated, 5 downregulated) that were significantly DE in eyes (log2fold change < |2|) (Table S7), including those downregulated in eyes: *DhpD1* and *sepia3* (pterins), as well as *cinnabar2* (ommochrome). For significant KEGG results, see Table S9.

### Shared differential expression for both photic tissues

We identified 1,075 genes differentially expressed with thorax in both light organs and eyes (600 upregulated, 475 downregulated in photic tissues), including seven pigment genes (Figure 6). Six of these genes were more highly expressed in both LO and eyes (i.e., DE with, or absent from thorax) (*DhpD2, sepia1, ras, scarlet, white1, lightoid*). One pigment gene displayed a divergent expression pattern between the photic tissues: *cinnabar2* (ommochrome) was upregulated in LO but downregulated in eyes relative to thorax. In addition, six other pigment genes were significantly DE with a log2 fold change < |2| relative to thorax (*KFase1, KFase2, cinnabar2, Sptr, Aprt, orange*) (Table S7). GO enrichment of genes upregulated in both LO and eyes relative to thorax (“photic genes”) are described in Table S11. Among these “photic” genes were nine LO genes from Fallon et al. (2018) (Table S10).

To understand which genes were important specifically in LO, we identified genes that were differentially expressed with thorax only in LO (i.e., DE only between LO vs. TX, but not eye vs. TX). Similarly, to understand which genes were important specifically in eyes, we identified genes that were differentially expressed with thorax only in eyes (and not in LO). Of the 2,037 genes DE between LO and thorax, a total of 1,260 genes were upregulated in LO. Among these were 43 LO genes from Fallon et al. (2018), of which nine were also upregulated in eyes relative to thorax and 34 were upregulated in LO only (Table S10). This included luciferase (PPYR_00001) and luciferin sulfotransferase (PPYR_00003), whose products are key for bioluminescence. No pigment genes were upregulated only in LO at a log2fold change > 2, however, four pigment genes that were upregulated in LO only (log2fold change < |2|), including two pterins (*mal, rosy3*). Cardinal (ommochrome) was exclusively downregulated (log2fold change < |2|) in LO, as well as two pigment genes downregulated at log2fold change < -2. In eyes, three pigment genes were upregulated (log2fold change > |2|) and two pigment genes were downregulated (log2fold change < |2|) (Figure 6; Table S7). GO enrichment of differentially expressed genes only in LO or eyes is reported in Tables S12-13.

### Contrast 3: Activity

We investigated transcriptional changes that occur in photic tissues of active fireflies by comparing pooled LO and eyes during signaling (active at dusk) and rest (inactive in the morning). Compared with Contrast 1 (light color), fewer genes were DE with activity in LO and eyes. Between LO activity state, 568 genes were DE (active: 393 upregulated 175 downregulated) (Figure 7). Of these, *Aprt* and *rosy1* (outlier) were upregulated in active LO, whereas *mal* was upregulated when fireflies were inactive (log2 fold change > |1|). Pigment genes DE with log2 fold change < |1| were DhpD1 (upregulated in active) and lightoid (upregulated while inactive). Only *Cytochrome-P450* (PPYR_06980) from Fallon et al. (2018) was upregulated in inactive LO. Between activity states in the eye, we detected just 43 differentially expressed genes (in active: 32 upregulated, 11 downregulated). Expression of pigment genes remained stable across activity states in the eye, with no differential expression (Figure 7).

**Figure 7.**
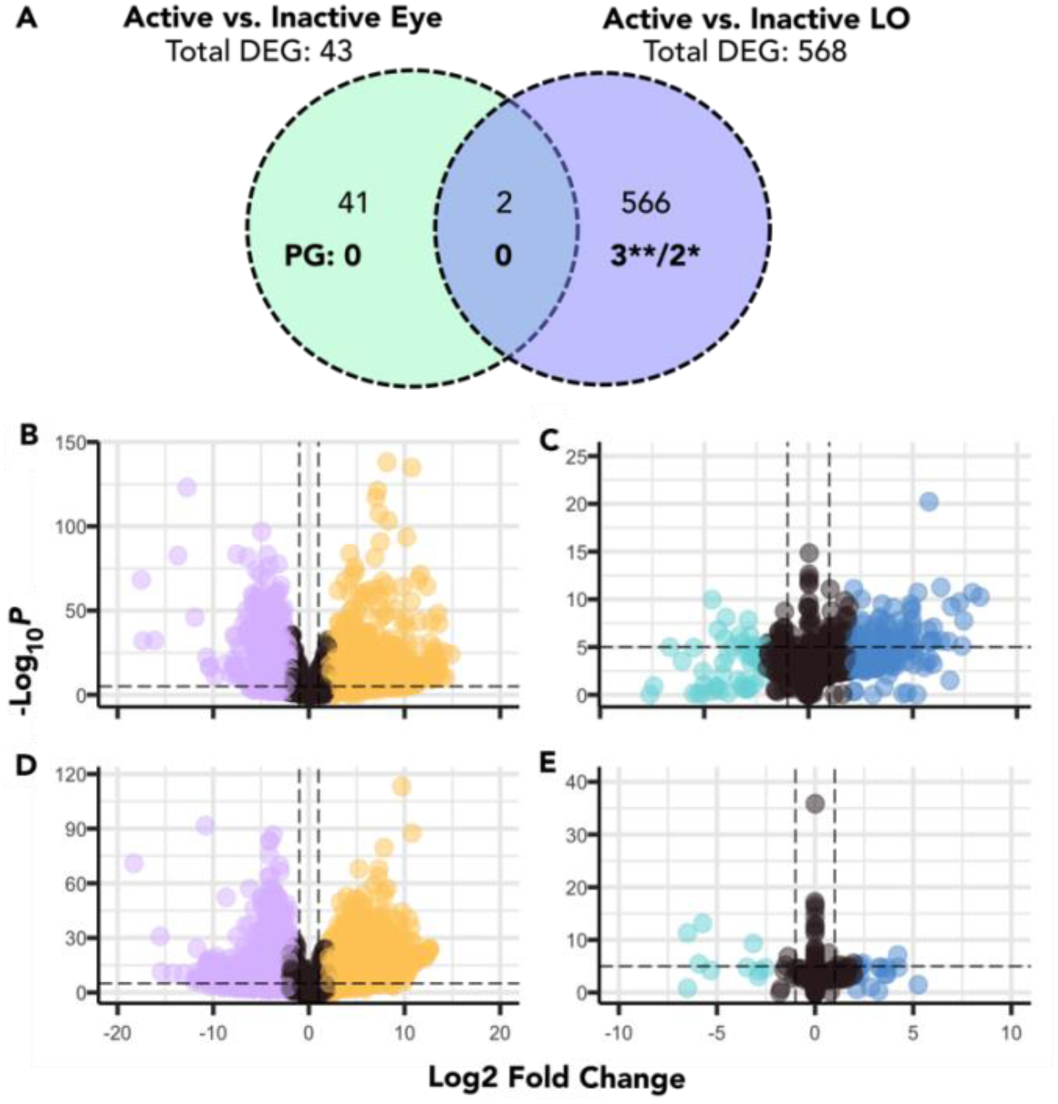
**A:** Differential gene expression in light organs and eyes between active and inactive fireflies. Of 43 genes differentially expressed between active and inactive eyes, none were pigment genes. Overall, 46 pigment genes were expressed in both states of LO and 47 in eyes. **Significant p-adjusted (BH) and log2fold change > |1|, *Significant p-adjusted (BH) DEG. Comparison of gene expression between **B:** LO vs. TX (active), **C:** Active vs. inactive LO, **D:** Eye vs. TX (active), **E:** Active vs. inactive eye. Dashed lines indicate thresholds used for p-value (horizontal) and log2 fold change (vertical) for Contrasts 2 and 3. Plots made with R package EnhancedVolcano (1.20.0).

To identify candidate genes associated with the hue change in LO, we identified genes that were upregulated in active LO (using both thorax and activity contrasts). Of these 150 genes, none were pigment or LO genes from Fallon et al. (2018); however, as these genes could be otherwise relevant to signaling in LO, the list is reported in Table S14.

### Head shield pigments (aposematic signal)

Overall, in the head shield (pronotum) 45 pigment genes were expressed >3 TPM (Table S6). Notably, *sepia1* (pterin) and *scarlet* (ABC transporter) were not expressed (<3 TPM) in HS, though differentially expressed with thorax in both LO and eyes. A total of 1,133 genes were in the 90^th^ percentile of highest expression (HE) in the headshield, including 10 pigment genes (Table S15). These included two pterins (*rosy1, rosy 3*), two ommochromes (*vermilion, cinnabar2*) and one transporter gene (*white1*). Of these, *white1* was also upregulated in both LO and eyes versus thorax and *cinnabar*2 was only upregulated in LO (not eyes) vs. thorax (log2fold change <2).

### Network analysis of light organ samples

A total of 12,572 genes were used to build a signed network (blockwiseModules function, R with power=14 and default parameters, yielding 40 different co-expression modules ranging in size from 33 to 1,829 genes (x̄= 314 ± 372 genes) (Tables S16-17). In 23 of these modules, we identified at least one pigment gene. There were 62 hubs significantly (adjusted p-value < 0.05) associated with activity state across 10 modules; none were pigment genes (Table S18). No hubs were associated with light color. Two modules had marginally significant (p<0.05, p-adjusted>0.05) associations with light color (Figure S9).

### Modules associated with light organ activity state

Three modules were significantly associated (p-adjusted < 0.05) with activity state: The genes in module M-24 (“pink”) had a significantly higher expression in inactive light organs, while the genes in modules M-12 (“green”) and M-33 (“skyblue3”) were significantly higher expressed in active light organs (M=12 had borderline significance, with p-adjusted= 0.0551); Table 2; Figure 8). We considered nine other modules with marginal significance of interest, with five modules more highly expressed in active light organs and four more highly expressed in inactive light organs (Table 2). For our analysis of activity state, we focused primarily on the three modules with greatest statistical association with activity state: M-33, M-12, M-24 (additional information in Supplemental Results). Module characterization (e.g., hub genes, enrichment, and GO terms) are reported in Supplemental Tables S18-20.

**Table 2.**
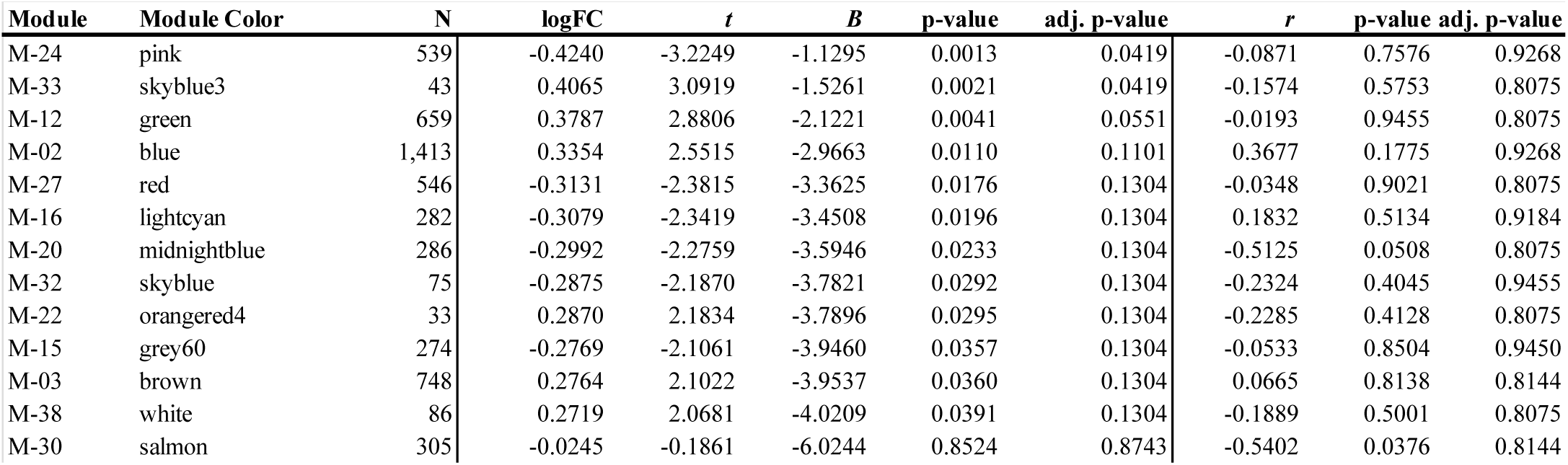
Modules significantly associated with the light organ phenotypes activity state and/or emitted light color.

| Module | Module Color | N | logFC | *t* | *B* | p-value | adj. p-value | *r* | p-value | adj. p-value |
| --- | --- | --- | --- | --- | --- | --- | --- | --- | --- | --- |
| M-24 | pink | 539 | -0.4240 | -3.2249 | -1.1295 | 0.0013 | 0.0419 | -0.0871 | 0.7576 | 0.9268 |
| M-33 | skyblue3 | 43 | 0.4065 | 3.0919 | -1.5261 | 0.0021 | 0.0419 | -0.1574 | 0.5753 | 0.8075 |
| M-12 | green | 659 | 0.3787 | 2.8806 | -2.1221 | 0.0041 | 0.0551 | -0.0193 | 0.9455 | 0.8075 |
| M-02 | blue | 1,413 | 0.3354 | 2.5515 | -2.9663 | 0.0110 | 0.1101 | 0.3677 | 0.1775 | 0.9268 |
| M-27 | red | 546 | -0.3131 | -2.3815 | -3.3625 | 0.0176 | 0.1304 | -0.0348 | 0.9021 | 0.8075 |
| M-16 | lightcyan | 282 | -0.3079 | -2.3419 | -3.4508 | 0.0196 | 0.1304 | 0.1832 | 0.5134 | 0.9184 |
| M-20 | midnightblue | 286 | -0.2992 | -2.2759 | -3.5946 | 0.0233 | 0.1304 | -0.5125 | 0.0508 | 0.8075 |
| M-32 | skyblue | 75 | -0.2875 | -2.1870 | -3.7821 | 0.0292 | 0.1304 | -0.2324 | 0.4045 | 0.9455 |
| M-22 | orangered4 | 33 | 0.2870 | 2.1834 | -3.7896 | 0.0295 | 0.1304 | -0.2285 | 0.4128 | 0.8075 |
| M-15 | grey60 | 274 | -0.2769 | -2.1061 | -3.9460 | 0.0357 | 0.1304 | -0.0533 | 0.8504 | 0.9450 |
| M-03 | brown | 748 | 0.2764 | 2.1022 | -3.9537 | 0.0360 | 0.1304 | 0.0665 | 0.8138 | 0.8144 |
| M-38 | white | 86 | 0.2719 | 2.0681 | -4.0209 | 0.0391 | 0.1304 | -0.1889 | 0.5001 | 0.8075 |
| M-30 | salmon | 305 | -0.0245 | -0.1861 | -6.0244 | 0.8524 | 0.8743 | -0.5402 | 0.0376 | 0.8144 |

**Figure 8.**
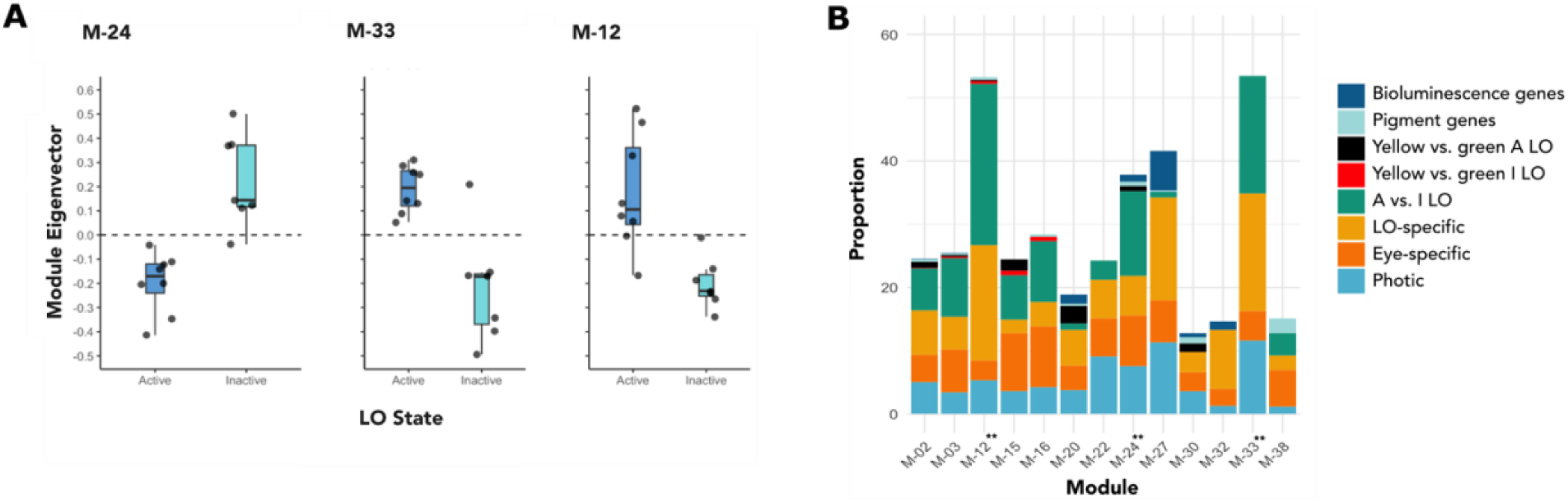
**A:** Gene network modules with altered expression patterns that are significantly associated with light organ activity. The 659 genes in M-12 (“green”) and the 43 genes in M-33 (“skyblue3”) were more highly expressed in active light organs; the 539 genes in M-24 (“pink”) were more highly expressed in inactive light organs. **B:** Proportion of pigment and bioluminescence genes, as well as DEG, within significant modules (** denotes highly significant, p<0.05 and p-adjusted <0.05).

Two of these modules (“green” and “skyblue3”) are associated with active LO. The “green” module includes pigment genes (pterins *DhpD1, sepia1*, and *clot*) in addition to genes associated with circadian rhythm and light perception, such as *arrestin domain-containing protein 17-like* (PPYR_02715) (Table S17). We identified 13 “hub genes” (Table S18) in addition to DE genes in the “green” module that distinguish yellow and green light organs (n=4) as well as yellow and green active eyes (n=2). The “skyblue3” module was small with only 43 associated genes, of which eight were hubs (Table S18), including *aristaless-related homeobox protein-like | retinal homeobox protein Rx1-like* (PPYR_00572), a gene involved in vertebrate eye development (Nelson et al, 2009). This module also contained *phenoloxidase-activating factor 1-like* (PPYR_14004) and *CLIP domain-containing serine protease 2-like|phenoloxidase-activating factor 3-like* (PPYR_01625) which activate melanogenesis in the immune response. There were 16 genes in “skyblue3” that were annotated as “uncharacterized protein” or lacked annotation, signifying specific-specific function. Both “green” and “skyblue3” modules were significantly enriched for genes DE with activity (“green” FDR=1.04E-84, “skyblue3” FDR=2.95E-03) LO-specific genes (“green” FDR=2.63E-35 and “skyblue3” FDR= 9.06E-03).

The “pink” module included transcripts upregulated in inactive LO. Four of these are pigment genes (ommochrome: *cardinal*, ABC transporter: *scarlet*, granule: *lightoid, pink*). *Pink* is part of the biogenesis of lysosome-related organelles complex (BLOC1-4), which is involved with the formation of pigment granules via intracellular trafficking (Falcón-Pérez et al., 2006). Other members of the BLOC family were observed: *ras-related protein Rab-32|ras-related protein Rab-38* (PPYR_05587), encoded by BLOC-3 (Gerondopoulos et al., 2012), and *biogenesis of lysosome-related organelles complex 1 subunit 4* (PPYR_10058). As with modules “green” and “skyblue3,” we observed genes related to phototransduction, including four copies of *retinol-binding protein pinta-like* (Tables S17). We additionally noted genes that play a role in eye development and/or maintence: *homeobox protein OTX1-like|homeobox protein OTX2-like* (PPYR_13979), *chaoptin-like* (PPYR_14596), *rab11 family-interacting protein 2* (PPYR_05444), and *ras-related protein Rab-28-like* (PPYR_12570).

Along with signal reception, there were six bioluminescence-related genes in module “pink” (Table S17), including *luciferin-sulfotransferase* (PPYR_00003) which catalyzes luciferin synthesis, a key component of the bioluminescent reaction. This module also included genes associated with photoreception, both of which were upregulated only in eyes relative to thorax and differentially expressed between light organ activity state (PPYR_12570, PPYR_12713). We also identified DEGs that separate yellow and green light organs (n=4) and eyes (n=8) (Table S17), as well as 19 hubs (Table S18). Module “pink” was signficantly enriched for photic genes (FDR= 1.25E-02) and genes DE with activity (FDR= 1.30E-16).

## Discussion

Our results show that all 47 pigment genes in the ommochrome and pterin biosynthesis pathways were expressed (>3 TPM) in eyes, and 46 pigment genes were expressed in the LO of *P. pyralis* fireflies (Table S6). Therefore, the production, transport and/or storage of the colored pigments generated by these genes could indeed filter the light generated by luciferase in LO. We did not identify any differentially expressed (DE) pigment genes between the LO of *P. pyralis* with “yellow” (565-568 nm) and “green” (560-562 nm) light color, contrary to our initial expectation. In eyes, only one pigment gene, *sepia3,* was upregulated in fireflies with yellower light. Thus, detecting subtle differences in gene expression relevant to this variation may require larger numbers of replicates to achieve greater statical power. Nevertheless, our tissue and activity contrasts yielded important insights.

We identified the differential expression of genes in ommochrome and pterin biosynthesis pathways in LO, including a set of genes also expressed in eyes, when compared with the thorax, a non-photic reference. During their active state, *P. pyralis* fireflies expressed 15 “photic” genes. Interestingly, 14 of these pigment genes shared the same directionality (i.e., upregulated or downregulated relative to thorax), which suggests the same types of pigments in tissues could be synthesized for use in light signaling. Importantly, we identified five pigment genes involved in consecutive steps of the ommochromes pathway: *KFase1, KFase2, cinnabar2*, as well ABC transporters *white1* and *scarlet*. Following the conversion of formyl kynurenine into kynerunine by KFase, the enzyme encoded by *cinnabar* converts LK into the yellow pigment 3-hydroxy kynurenine (3-OHK), which can either remain in the cytoplasm to impart coloration (Llandres et al., 2013) or be imported into ommochrome granules by *white* and *scarlet*. The expression of these genes in both LO and eyes 3-OHK is synthesized and relocated inside the granules, where it gets converted into xanthommatin (yellow-brown) non-enzymatically via dimerization of 3-OHK (Figon et al., 2020). Reduction of xanthommatin results in dihydroxanthommatin (red), which can be converted into ommins (violet) via either non-enzymatic condensation of xanthommatin with a sulfur-containing derivative of methionine/cysteine (Figon et al., 2020) or cardinal (Xu et al., 2020).

As many steps of the pathway occur non-enzymatically, we were unable to resolve the extent of ommochrome synthesis in LO through gene expression alone and present plausible outcomes based on these patterns. We found *cardinal* was lowly expressed in LO (mean TPM±SD 6.20±4.54) compared with eyes (mean TPM±SD 600.88±206.48), and downregulated relative to thorax (Contrast 2). Decreased cardinal activity and/or lack of nonenzymatic conversion of 3-OHK into xanthommatin could correspond with high levels of 3-OHK, which provides yellow coloration in crab spiders (Riou & Christidés, 2010) and *Heliconius* butterflies (Finkbeiner et al., 2017), with the latter using 3-OHK in both eyes and wings for mate recognition. Cardinal knockouts in *Bombyx mori* resulted in red eyes, as opposed to their dark coloration in wildtype (Osanai-Futahashi et al., 2016), as well as the accumulation of 3-OHK in *Drosophila* (Harris et al., 2011; Shirai & Daimon, 2020). Alternatively, conversion of 3-OHK to xanthommatin through non-enzymatic reactions could result in production of yellow-brown, red, and/or violet ommochromes, including those implicated in reversible color change due to shifts in redox state (Futashi et al., 2012). Overall, our results suggest that 3-OHK is incorporated into granules, where subsequent conversion can occur.

In the pterin pathway, upregulated “photic” genes included *DhpD2* and *sepia1*. In addition, three copies of *sepia* (*sepia2, sepia3, sepia5*) were expressed in all four tissues (LO, eyes, thorax, HS) (Table S6), raising the possibilty that their colored products could influence light filtering and tuning between LO and eyes. Though we cannot verify these transcripts represent duplicated genes, this observation suggests some pigment gene families have duplicated; how their products differ in enzymatic activity and generation of colored products remains to be investigated. The synthesis of orange pigments requires *DhpD*, which encodes dihydropterin deaminase that transforms 7,8-dihydropterin into 7,8-dihydrolumazine (yellow) that undergoes condensation into aurodrosopterin (orange). Encoded by sepia, pyrimidodiazepine synthase generates several pigments, including the yellow sepiapterin (Figure 3). Conversion of 6-pyruvol-tetrahydropterin into pyrimidodiazepine (Kim et al., 2006) requires thioredoxin reductase, the product of *clot* (Giordano et al., 2003; Wiederrecht et al., 1984). Finally, pyrimidodiazepine condesenses into either aurodrosopterin or drosopterin or its isoform, isodrosopterin (red) (Schwinck, 1975; Rokos and Pfleiderer 1975; Ayling et al., 2012; Andrade & Carneiro, 2021). While *clot* was not differentially expresed with thorax in either LO or eyes, it was expressed at similar levels (LO: mean TPM±SD=21.87±10.88, eyes: 19.25±3.69), suggesting it functions with *sepia* to produce orange and/or red pterins in photic tissues. This is supported by our preliminary pigment data from single pooled samples (N=2) (Table S22) because the relative abundance of 7,8-dihydrolumazine was more than double in active yellow (6.72%) and yellow-green (8.97%) LO compared to green LO (2.89%); however, since our analysis did not standardize by input tissue, these differences could be explained by variation in firefly body size. Further, Welch two-sample t-test (Supplement) between active yellow and green LO revealed no difference in the relative abundance of pterin substrates (p=0.9998) and clustering analysis indicated high similarity between yellow, green, and yellow-green LO (Figure S11), suggesting light color variation is not explained by differences in pterin substrates. In parallel, 7,8-dihydrolumazine was more abundant in yellow (77.26%) and yellow-green (66.77%) eyes compared to green eyes (28.48%), providing a possible mechanism of tuning between light emission spectra and visual sensitivity. We identified low levels of drosopterin but no aurodrosopterin in LO, suggesting that 7,8-dihydrolumazine is the most abundant pterin and could contribute yellow coloration (Table S22). This observation must be verified through increased sampling and inclusion of standards. Metcalf (1943) described “lampyrine,” a pink pigment found across body tissues only in Lampyridae (Coleoptera) that shared some characteristics with pterins. This raises the possibility that fireflies may synthesize unique, lineage-specific pigments that contribute to color, yet remain undetected.

Two pterins were uniquely upregulated in LO relative to thorax: rosy3 and mal, suggesting they could perform a specific function in LO. The *rosy* locus encodes the enzyme xanthin dehydrogenase (XDH), which is involved in several enzymatic reactions, such as the transformation of pterin (colorless) into isoxanthopterin (colorless), 7,8 dihydropterin into the precursor of xanthopterin: H2-xanthopterin (yellow), and xanthopterin (yellow) into leucopterin (colorless) (Ferré, 2024), as well as the production of uric acid (Hilliker et al., 1992; Rizki & Rizki, 1962), which is abundant in LO. Notably, *rosy3* was more highly expressed in the active green LO (mean TPM±SD=11,177.25±6,400.87) than active yellow LO (mean ±SD=6,456±1,477.64), though *rosy3* was not significantly differentially expressed between active yellow and green LO. Our PCA analysis reveals leucopterin clusters with all three LO samples (Figure S11), suggesting *rosy* activity could also underlie production of leucopterin in LO. There is a direct relationship between the number of copies of *rosy* and XDH activity (Hubby and Forrest, 1960; Glassman et al., 1962; Grell, 1962), and at least three copies (*rosy1, rosy2, rosy3*) were expressed in all *P. pyralis* tissues (Table S6). *Mal* encodes molybdenum cofactor sulfotransferase, which is required for *rosy* (pterin) activity (Finnerty et al., 1979; Browder et al., 1992). *Drosophila* without *rosy* and *mal* activity had brown eyes due to a lack of purines and pterins (Hubby & Forrest, 1959), indicating their role in red coloration (Reaume et al., 1989). That our computational analysis identified these interacting partners expressed in LO clarifies how enzyme pathways are conserved and deployed in nonmodel species, though functional analysis is needed to reveal the ways in which pigment synthesis contributes to adaptive coloration.

In *Photinus* fireflies, the head shield (HS) has conspicuous pink coloration with black contrast and is used as an aposematic signal, advertising toxicity (lucibufagens) to predators (Day, 2011). To identify the pigment genes underlying this static pink coloration, we identified genes among 90^th^ percentile of highly expressed (HE) genes in HS. Insects are known to redeploy pigment expression networks across tissues and developmental stages, resulting in intraspecific variation (Wittkop et al., 2002; Vargas-Lowman et al., 2019) and the evolution of novel traits involved with coloration (Reed & Nagy, 2005; Ferguson & Jiggins, 2009; Monteiro, 2012; Martin et al., 2014). Overall, 44 pigment genes were expressed in HS, LO, and eye. To determine if pigments used for interspecific communication could also be used in LO for filtering emitted light, we compared our differential expression lists with genes HE in the HS.

Among these shared pigment genes were *rosy3* and *Prat2*, which were “LO-specific” (upregulated only in LO vs. thorax), as well as *cinnabar2*; while *cinnabar2* was DE in both LO and eyes relative to thorax, it was upregulated only in LO. This suggests that HS produces 3-OHK, which may contribute yellow coloration in both LO and HS. Importantly, *white1* was HE in HS and “photic,” indicating its relevance to the three tissues (LO, eyes, HS) that use pigments for signaling. As *scarlet* was not expressed in HS, the high expression of *white1* implies its use in the pterin pathway with an unknown binding partner, as we did not recover a *brown* ortholog possibly due to divergence (Grubbs et al., 2015). Lack of *scarlet* expression in HS indicates that 3-OHK remains in the cytoplasm without further modification as a yellow pigment. Interestingly, *scarlet* and *sepia1* were expressed only in LO and eyes, implying they could play a role in light signaling.

The transition from a resting to signaling firefly involves diel changes in physiology, such as the hue change observed in *P. pyralis* prior to the onset of activity at twilight. Interestingly, a much greater transcriptional change occurred between activity states in LO compared to eyes, indicating there is a heightened biological response in LO in preparation for signaling. In contrast, no pigment genes were differentially expressed with activity in eyes. As screening pigments are integral for vision and protect eyes from UV light, pigment genes in eyes may be consituently expressed. In LO, 46 pigment genes were expressed with no difference across states (Table S6), with the exception of three that were upregulated in active LO: *rosy1* (LO outlier, Figure S8), *DhpD1*, and *Aprt*, and two downregulated: *mal* and *lightoid*. *DhpD1* was the most highly expressed *DhpD* copy across tissues, and *DhpD* generates yellow 7,8-dihydrolumazine. This was supported by our preliminary LC-MS analysis of pterin substrates in *P. pyralis* LO, eyes, and HS (Supplement), as we identified yellow 7,8-dihydrolumazine (among other pterins) in both active and inactive LO and eyes (Figure S11; Table S22). Overall, the different tissue extracts displayed a unique pterin pigment profile (Figure S11; Table S22), and a PCA clustering analysis suggests there may be differences between activity state (Figure S11).

In *Photinus* eyes, an unknown type of magenta-colored screening pigment (proposed to be a pterin) has been implicated in tuning visual sensitivity (Cronin et al., 2000). We cannot identify which of the screening pigments in our study would extract as a magenta pigment, and whether our “unknown” could be this pigment. Few pigment genes were differentially expressed with activity in *P. pyralis* LO, so the hue change is likely mediated through additional mechanisms. Specifically, this rapid change could arrise from posttranslational processes such as changes to the intracellular environment like pH or redox state. We identified significantly differentially expressed genes (DEG) related to cellular pH, which could be connected to changes in redox state due to homeostasis and/or within organelles that could potentially alter phenotype. For example, xanthommatin (yellow-brown) is reduced into dihydroxanthommatin (red) under acidic conditions in male dragonflies (Futahashi et al., 2012). Several GO terms related to shifts in pH (i.e., proton transmembrane transport) were recovered among genes upregulated in LO-specific genes (Table S12), as well as the “lysosome” KEGG pathway for this same contrast, with higher expression of genes encoding acidification regulators (Table S8, Supplement). In M-27 “Red,” associated with inactive LO (p= 0.018, p-adj= 0.130) and also enriched for LO genes from Fallon et al. (2018), we observed GO terms vacuolar acidification, pH reduction, and response to alkaline pH, suggesting an association between genes related to pH and signaling, such as luciferase (Supplement).

Changing the intracellular conditions, such as pH, may shift which reactions are favored or increase rates of non-enzymatic condensation; for example, the pterin substrate 7,8-dihydrolumazine undergoes non-enzymatic condensation reactions with pyramidodiazepine (PDA) to produce aurodrosopterin at low pH (Yim et al., 1993; Kim et al., 2009). Though reversible, binding to proteins or metals can stablize the pigment by preventing interaction with other molecules despite changes in pH (Figon et al., 2021). Hence, processes outside the direct synthesis pathways could potentially change pigment pH and/or redox state, thereby influencing pigment phenotypes. These alternate pathways deserve greater attention from future functional assays to test new hypotheses generated by our computational genetic screens.

Given the interdependence and complexity of gene expression, we used network analysis to investigate patterns of co-expression associated with the phenotypes LO activity state and emitted light color, which revealed three modules signficantly associated with LO activity state. These network analyses validate and extend our DEG analyses with independent assessment of gene underlying change in activity state. Of the two modules correlated with active LO, M-33 (“skyblue3”) and M-12 (“green”), pigment genes were present only in M-12, with three genes from the pterin pathway (*DhpD1*, *sepia1*, and *clot*). This supports the importance of *DhpD* identified in our gene expression and pigment analyses. Interestingly, *DhpD1* was downregulated in both LO and eyes relative to thorax (Contrast 2), but upregulated in active LO (Contrast 3), whereas *sepia1* was “photic” (Contrast 2). As the catalytic activity of *sepia* requires *clot*, the coordinated expression of genes involved in biosynthesis of yellow, orange, and/or red pterins with active LO suggests a link between signaling state and LO pigmentation. Other copies of these genes (*DhpD2, sepia3*) were present in M-13 (“greenyellow”), which was not associated with light organ state, suggesting that different copies my vary in function. Both, the elevated copy number (Table 1, Figure S5) and expression divergence (Table S7) of *DhpD* and *sepia* copies imply that these genes have undergone duplication and neofunctionalization, a furthering this as a direction for future studies.

Genes in M-27 (“pink”) were more highly expressed in inactive LO and included four pigment genes: *scarlet*, *cardinal*, *lightoid*, and *pink*. The co-expression of two genes in the ommochrome pathway (*scarlet*, *cardinal)*, strengthen our confidence that yellow, orange, red, and/or violet ommochromes are produced in LO regardless of activity, and could contribute to tuning light signals in both LO and eyes. Genes involved in the biogenesis of pigment granules emerged from this module. Pigments are considered lysosomal-related organelles because they share characteristics with lysosomes, including acidified lumens (Dell’Angelica et al., 2000; Ohkuma & Poole, 1978) and generation (Luzio et al., 2014). *Pink* encodes a subunit of the biogenesis of lysosome-related organelles complex (BLOC) which are implicated in the formation of screening pigments (Falcón-Pérez et al., 2006) and are critical for body coloration. For example, *Drosophila pink* mutants displayed a decline in the abundance of red and brown pigments compared with wildtype (Falcón-Pérez et al., 2006). The decrease in these pigments was compounded by loss of function in the rab GTPase *lightoid* (Ma et al., 2004; Falcón-Pérez et al., 2006). Interestingly, *Lightoid* interacts with pigment granules (Fujikawa et al., 2002; Satoh et al., 2008) and was upregulated in both photic tissues (Contrast 2) as well as downregulated in active LO (Contrast 3, log2 fold change=-0.59114), with expression in HS (TPM=686). *Lightoid* plays a key role in the pigment migration, which insects use as like a pupil to enhance vision (Stavenga & Kuiper, 1977). In *Drosophila* eyes, *lightoid* links granules to the myosinV motor for transport along actin filaments of the cytoskeleton (Satoh et al, 2008).

Fascinatingly, M-27 was enriched for “photic” genes upregulated in both LO and eyes relative to thorax (Table S19) and the “retinal metabolic process” GO term (Table S20), in addition to four copies of *retinol-binding protein pinta-like* (Table S17). *Pinta* is integral for vision in *Drosophila* and is expressed in retinal pigment cells (Wang & Montell, 2005), raising questions about why genes characteric of eyes are associated with inactive LO, and if this transcription has functional consequences. It is possible that genes related to circadian rhythm, which relies upon light perception (Wijnen et al., 2006; Yoshii et al., 2016) to mediate shifts between activity state. Importantly, also present in M-27 was *luciferin sulfotransferase* (PPYR_00003), as well as four other LO genes from Fallon et al. (2018), indicating that members of the ommochrome pathway and granule group are regulated alongside genes essential for light signaling. As pigment biosynthesis pathways are complex with interrelated components, these results enhanced our understanding of the processes that accompany signaling in fireflies.

As our second question we asked whether the differential expression of pigment genes could explain variation in light color between two population of *P. pyralis* fireflies with identical luciferases, resulting in differences in the abundance and/or types of pigments in LO. Our initial question asked whether the differential expression of pigment genes could explain variation in light color, which assumes that morphological changes arise from differences in the abundance and/or types of pigments in LO. In addition to natural intraspecific variation, the *size* and *location* of pigments within the cell also influence their filtering effect. For example, the fiddler crab *Uca panacea* undergoes a diel color change due to the dispersion of the pigment granules (Darniell, 2012). Studies of firefly granules in LO indicate that granules are located at the periphery of photocytes (Ghiradella & Schmidt, 2004). At night their dark pigments aggregate, increasing light reflection so their carapace appears white. During the day, the crabs turn dark due to dispersion, or expansion of these light-absorbing granules. This phenomenon has been widely studied in fish and crustaceans, where it occurs in both eyes and body tissues; similar physiological color changes that affect orange or red body coloration have been observed in krill (Auerswald et al., 2008) and gobies (Svensson et al., 2005). Components involved in pigment migration are conserved; pterins found across animals (Andrade & Carneiro, 2021) and ommochromes are distributed among invertebrates (Needham, 1974), whereas signals involved in pigment migration, such as pigment dispersing hormone are common in arthropods (Meelkop et al., 2011; Rao & Riehm, 1989). Therefore, a similar mechanism could be used to generate the pigment filter in fireflies, possibly alongside changes to molecular structure such as redox state. While differential expression of genes in pigment biosynthesis pathways cannot explain intraspecific variation in light color, we provided evidence that ommochromes and pterins could be involved in tuning light signals, as well as candidate pigment genes (e.g., *DhpD*). Using whole transcriptome computational analysis, we identified differential expression patterns that could support alternative explanations that may be more complex than differences in pigment synthesis and extend to the modification of pigment molecules and/or translocation of their granules. As physiological color change is rapid and reversible, this effect of pigment migration could generate the hue change in *Photinus* LO and be a valuable focus for future study of covariation between signaling spectrum and habitat change through time.

**Supplemental materials:** https://github.com/mspopecki/Gene-expression-Photinus-pyralis.git

## Acknowledgments

Rebekah Rogers and Margot Popecki are funded by NIH NIGMS MIRA R35 GM133376 to RL Rogers. We are extremely grateful for our insightful conversations with Dr. Florent Figon, Dr. Marley Brimberry, Dr. Bill Lanzilotta, and Dr. Jonathan Eggenschwiler. Thank you, Dr. Karolina Heyduk, Dr. Daniel Shaw, Dr. Gaelen Burke, Dr. Kaixiong Ye, and Dr. Doug Menke for your input on experimental analyses, and Nicole Steel for molecular work. Funded by the University of Georgia (UGA) Graduate School, UGA Department of Genetics, and UGA Museum of Natural History. Haley DeLoach cheerfully provided her assistance with field collections, which would not have been possible without generosity of The Pastures of Rose Creek. This paper is dedicated to the memory of Will Breedlove, may his joy forever be remembered in the “yellow” fireflies at twilight.

