## Supplement for "The role of pigments in light color variation of the firefly *Photinus pyralis*"

### Pigment Analysis

In parallel to our approach for RNA extraction, we dissected eyes, light organs, and head shield tissues from ten additional *P. pyralis* specimens (six active, four inactive). Due to the limited number of flash-frozen fireflies at the green and yellow end of the light spectrum, we pooled the pigment extracts from two fireflies (each) with similar light emission spectra, for a total of 3 samples for both active light organs and active eyes for pigment analysis: (1) emitting green ($\bar{x}$=561.46± 0.067 nm), intermediate yellow-green ($\bar{x}$=563.08±0.311 nm), and yellow ($\bar{x}$=565.30±0.306 nm) light color. In addition, we prepared one inactive sample (2 fireflies) emitting yellow light color ($\bar{x}$= 565.080± 0.857 nm). For comparison of the relative abundance of pterin substrates in LO and eyes, we also pooled the head shields from three active *P. pyralis* specimens to ensure sufficient pigment concentration for analysis.

### Pigment extraction from firefly tissues

Following Rutowski et al. (2005) for extraction of pterin pigments, we added 400 µL of 1% $NH_4OH$ (in water) to each tissue sample. For extraction of ommochromes pigments, we used 400 µL of 0.5% HCL in methanol (Llandres et al., 2013; Fabricant et al., 2013) before grinding the tissues with plastic pestles and an electric hand-held homogenizer until fully disrupted. Extracts were incubated in the dark on a shaker at low speed at 4°C for 48 hours. Extracts were then centrifuged with a Nanosep 0.2 µm BioInert membrane spin column (Pall Corporation) at 5,000 rcf for 15 minutes.

### Measuring the absorbance spectra of pigment extracts with UV-vis

To determine if pigments in the light organ could potentially shift the light spectrum of the bioluminescent reaction, we measured the absorbance spectra of our total extracts across the UV and visible spectrum (UV-VIS 325-1000 nm) with a UV-Vis (OlisWorks HP 8453), using the extract medium as a blank. We analyzed the spectra with OlisWorks (v1.8.17063.294) and compared the peak absorbance of our samples with the published absorbance spectra of ommochromes and pterins (Figon & Casas, 2021; Roca-Sanjuán et al., 2014; Chen et al., 2007). After UV-vis, the remaining pterin extracts were frozen at -30°C until liquid chromatography with mass spectroscopy (LC-MS).

### Liquid chromatography-mass spectrometry (LC-MS)

The pterin pigment extracts were analyzed on a Thermo Vanquish (ThermoFisher Scientific) connected to a Thermo QExactive HF orbitrap mass spectrometer (ThermoFisher Scientific). A separation was carried out on a Supel Carbon LC; (Supelco; 2.1x100mm; 2.7 µm) with a separation condition as follows at 0.3m:/min: From 0-3 minutes the buffer was held at 90% acetonitrile in 0.1% formic acid, dropped to 50% acetonitrile and held for 8 minutes, and increased from 50% acetonitrile to 90% acetonitrile for another 12 minutes to equilibrate the column for the next run. The first 10 minutes of the separation was fed into the mass spectrometer for analysis while the equilibration was diverted to waste. A data-dependent program was used for acquisition where the precursor ion scan was acquired at 120k resolution followed by top-down fragmentation of high-to-low intensity m/zs by stepped HCD at 30k resolution. To test for the presence of pigments in the light organs, eyes, and the headshield, we used the MH+ values and the respective retention times (RT) of known compounds in the guanine-derived pterin pathway.

**UV-vis and LC-MS Pigment data**

We obtained the peak absorbance of pterin and ommochromes pigment extracts with UV-vis (Figure S10) and searched for 17 compounds from the pterin pathway in *P. pyralis* fireflies with LC-MS (Tables S21-22, Figure S11). Given that we only had  a single pooled (two specimens) sample each for active light organs and inactive light organs (and the corresponding pooled eye samples from the same fireflies) and no replicates, we could make limited comparisons. Pterin substrates with greatest abundance across all tissues were leucopterin, xanthopterin/isoxanthopterin, and 7,8 dihydrolumazine (though the order differs with tissue: Table S22). Unfortunately, our pigment analysis cannot distinguish between xanthopterin (yellow) and isoxanthopterin (colorless), nor red pigments isodrosopterin, drosopterin, and aurodrosopterin, so this remains to be tested with inclusion of standards that can distinguish isoforms and enantiomers. Low amounts of xanthommatin/isoxanthommatin and drosopterin/isodrosopterin/aurodrosopterin were detected in our preliminary analysis (Table S22). However, since we know only relative abundance (dependent on the abundance of other pigments) and not absolute amounts, we do not know how this translate to the actual pigment levels (e.g., substrate is present in each sample).

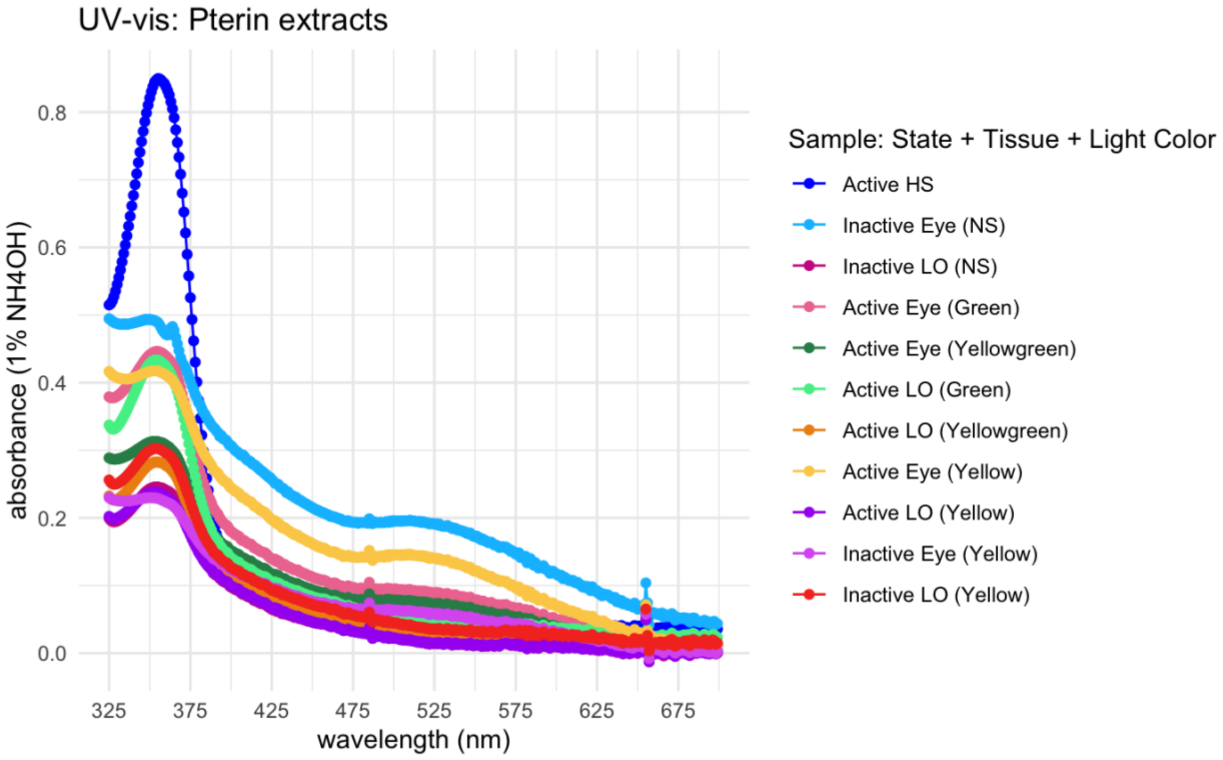


Figure S10: Absorbance measurements of pterin pigments from UV-vis. Blank was 1% $NH_4OH$. HS=head shield, NS=no spectra (unknown light color).

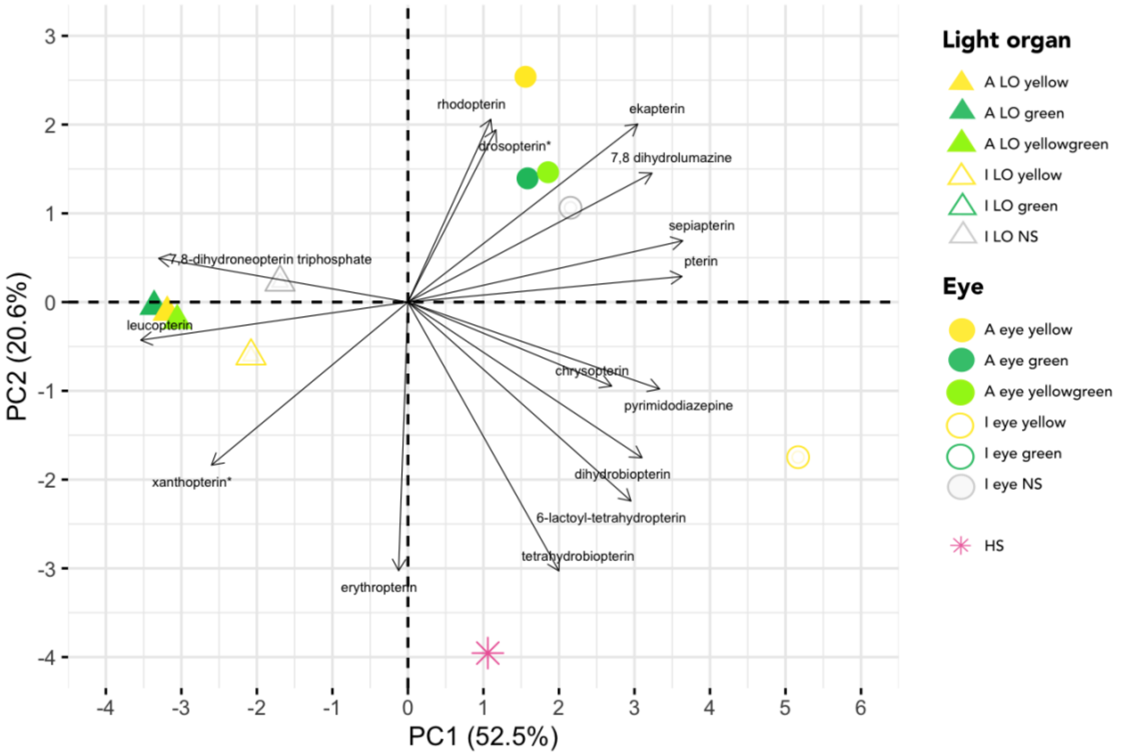


**Figure S11:** PCA Biplot of the 15 pterin substrates recovered in *Photinus pyralis* tissue samples. Pooled samples (N=2) are represented as points and pterin substrates by vectors. **Substrate has isoforms (i.e., same molecular weight).*

The main questions we aimed to answer with this study is whether screening pigments were present in the light organ, and if differences in these pigments could explain intraspecific variation in light color (i.e., green vs. yellow). Evidence from UV spectra suggests that pterins are present in both activity states of light organs, eyes, and head shield (Figure S10). All tissues displayed a peak around 370 nm, which is characteristic of pterins that absorb light <400 nm (Wijnen et al., 2007; Andrade & Carneiro, 2021). Though pterins also absorb light in the visible spectrum between 500-600 nm (Johnson & Fuller, 2015), we did not observe a second peak in this range. It is possible that eye samples display a shallow peak around 520 nm, but our data is inconclusive. This could be due to the low concentration of pterin extracts in all tissues. Thus, additional work is needed to characterize and quantify pterins in firefly light organs with the use of standards.

Our preliminary analysis suggests that the pigment composition of LO is largely driven by two pterin substrates: 7,8-dihydroneopterin triphosphate (possibly pale yellow to yellow) and leucopterin (colorless).

The pterin precursor 7,8-dihydroneopterin has antioxidant activity (Janmale et al., 2019) and is also involved with nitric oxide (NO) production (Crabtree et al., 2009). As NO is used to stimulate flashing in firefly LO (Aprille et al., 2004), this enzyme may have multiple roles related to bioluminescence and/or coloration. Leucopterin is a colorless pigment underlying the white coloration of *Pieris* butterfly wings (Wijnen et al., 2007) and is synthesized by xanthine dehydrogenase, encoded by *rosy*, which was expressed by three copies (*rosy1, rosy2, rosy3*), with particularly high expression of *rosy3* (mean TPM±SD= 9,272±4486.99) in LO.

There also appears to be some correlation between LO and xanthopterin (yellow) and/or isoxanthopterin (colorless), but our cannot distinguish between the isoforms. Based on vector position, there appears to be a negative correlation for relative abundances of drosopterin and its isoform, isodrosopterin (red), pterin (red), rhodopterin (brown), ekapterin (unknown color), 7,8-dihydrolumazine (yellow), and sepiapterin (yellow) in LO and eyes, with increased abundance in eyes (Figure S11). This implies red and yellow pterins are found at increased levels in eyes compared with LO, which aligns with the observed color differences between resting LO (ivory) and eyes (dark). We did observe low quantities of these six substrates in LO (Table S22), suggesting the pigments are present, though their filtering effect remains unclear. Repeating this analysis with increased replication and inclusion of standards could clarify whether colored pterins could contribute to tuning light color.

Overall, LO and eyes are characterized by distinct pigment profiles according to tissue (Figure S11). Aside from one sample (inactive yellow eye), tissues tend to group closely, indicating similarity. Light color does not appear to drive differences within tissue, which supports our finding that pigment gene expression does not explain variation in light color. In both LO and eyes, the inactive samples appear to group less closely with active, however more sampling is needed to determine if pigment substrates differ between activity states.

Wilcox (2021) showed species-specific patterns of UV-induced fluorescence across nine North American fireflies, noting high intensity in light organs; fluorescence can be used as a diagnostic to detect and characterize the presence pterins (Fabricant et al., 2013; Kayser 1985; Fox, 1976); future work could integrate measurements of fluorescence to verify this. Investigation of ommochromes in the firefly light organs would also clarify if both screening pigments are in LO and could influence emitted light color.

**Welch Two Sample t-test**

To determine if there were significant differences in the pterin substates between active samples with the most extreme "yellow" and "green" light color, we performed an unpaired t-test (Welch Two Sample t-test) in R (4.3.3). As input, we used the total percent relative abundance (m/z) of all pterin substrates for each sample. There were no significant differences between active "yellow" and "green" LO (p-value = 0.9998, df = 27.974), indicating that differences in the pterin substrates in active LO do not contribute to light color variation.

### Supplemental Figures

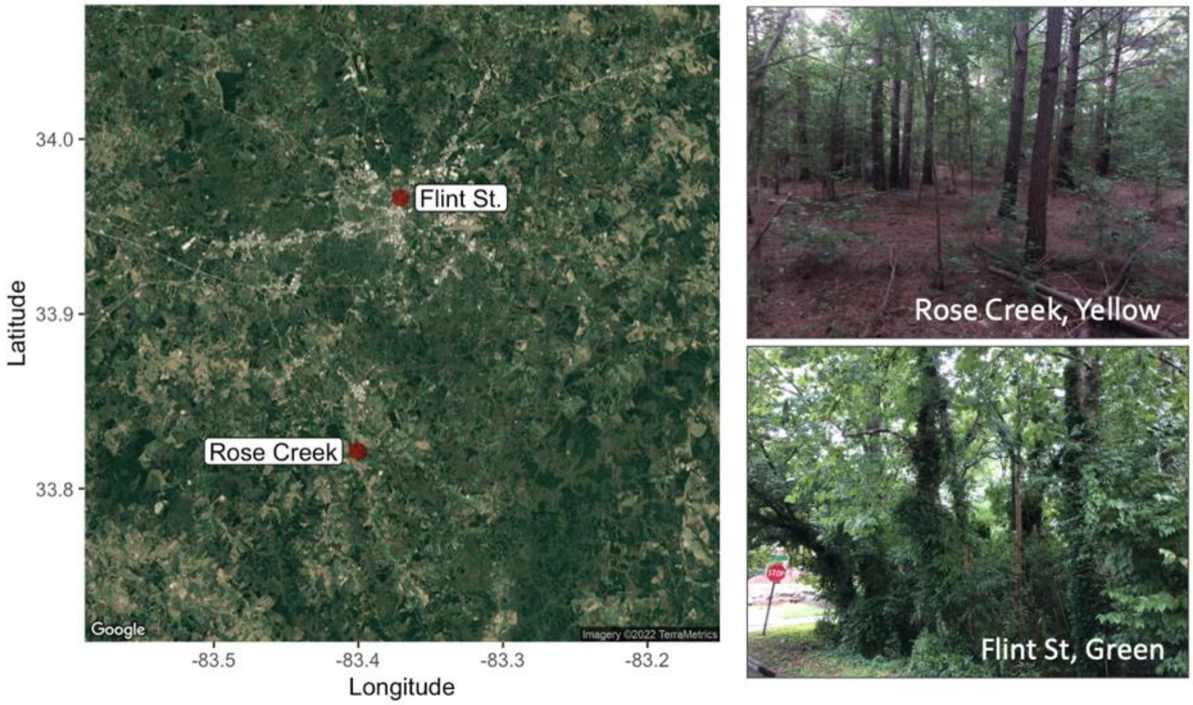


**Figure S1:** Sampling sites near Athens, Georgia (United States). Fireflies with greener and yellower light signals were collected at two sites: Flint St ("green") and Rose Creek ("yellow").

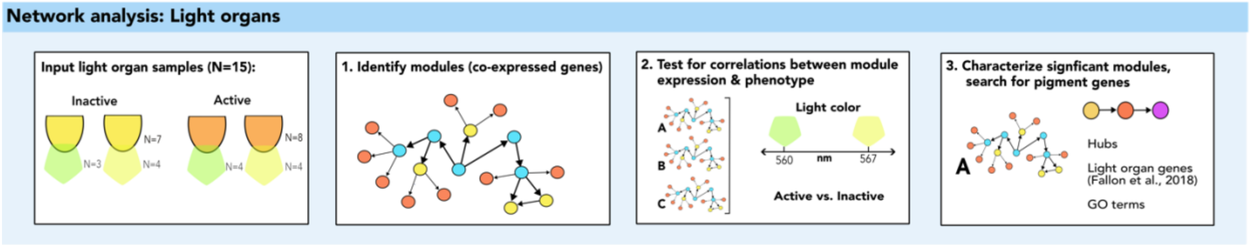


**Figure S2:** Network analysis with WCGNA. To understand patterns of gene expression in light organs, we identified sets of genes with correlated expression (modules) and then tested for correlations between modules and phenotype (activity state, emitted light color). We were particularly interested in modules that were significantly associated with a light organ phenotype and if any pigment genes were present.

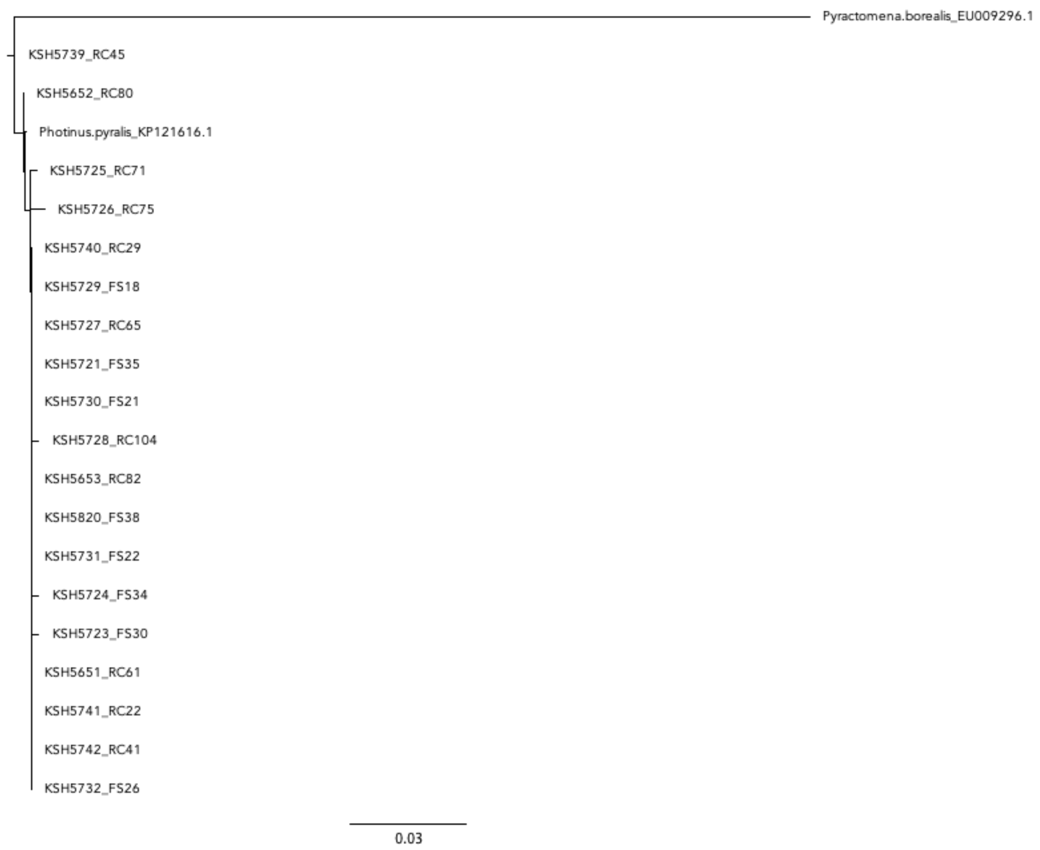


**Figure S3:** *Cytochrome oxidase subunit I* (COI) gene tree. To obtain the tree, partial COI DNA sequences from all samples were aligned in Geneious (2022.2.2) using MUSCLE. Included in the alignment were the COI sequences of the firefly *Pyractomena borealis* (accession EU009296.1) and *Photinus pyralis* (accession KP121616.1), obtained from GenBank. The alignment was trimmed to 543 bp as input to construct a neighbor joining tree, also within Geneious with the firefly *Pyractomena borealis* as outgroup. Our samples clustered in a clade with the confirmed *Photinus pyralis* sequence, verifying of field species identification for all specimens used in the transcriptome analysis.

```
Consensus                          -----------MEDAKNIKKGPAPFYPLEDGTAGEQLHKAMKRYALVPGTIAFTDAHIEV   49
TRINITY_GG_752_c0_g1_i1.p1_FS26A3  -----------.................................................   49
TRINITY_GG_731_c0_g1_i1.p1_RC41A3  -----------.................................................   49
TRINITY_GG_754_c0_g1_i1.p1_RC104I3 -----------.................................................   49
TRINITY_GG_807_c0_g1_i1.p1_RC29A3  -----------.................................................   49
TRINITY_GG_806_c0_g1_i1.p1_RC45A3  -----------.................................................   49
TRINITY_GG_751_c0_g1_i1.p1_FS35I3  -----------.................................................   49
TRINITY_GG_741_c0_g1_i1.p1_FS21A3  -----------.................................................   49
TRINITY_GG_796_c0_g1_i1.p1_FS22A3  -----------.................................................   49
TRINITY_GG_710_c0_g1_i1.p1_FS38I3  -----------.................................................   49
TRINITY_GG_734_c0_g1_i1.p1_RC65I3  -----------.................................................   49
TRINITY_GG_802_c0_g1_i1.p1_RC22A3  -----------.................................................   49
TRINITY_GG_758_c0_g1_i2.p1_RC75I3  -----------.................................................   49
TRINITY_GG_730_c0_g1_i1.p1_FS18A3  -----------.................................................   49
TRINITY_GG_756_c0_g1_i1.p1_FS30I3  -----------.................................................   49
TRINITY_GG_730_c0_g1_i2.p1_RC71I3  -----------.................................................   49

Consensus                          NITYAEYFEMSVRLAEAMKRYGLNTNHRIVVCSENSLQFFMPVLGALFIGVAVAPANDIY  109
TRINITY_GG_752_c0_g1_i1.p1_FS26A3  ............................................................  109
TRINITY_GG_731_c0_g1_i1.p1_RC41A3  ............................................................  109
TRINITY_GG_754_c0_g1_i1.p1_RC104I3 ............................................................  109
TRINITY_GG_807_c0_g1_i1.p1_RC29A3  ............................................................  109
TRINITY_GG_806_c0_g1_i1.p1_RC45A3  ............................................................  109
TRINITY_GG_751_c0_g1_i1.p1_FS35I3  ............................................................  109
TRINITY_GG_741_c0_g1_i1.p1_FS21A3  ............................................................  109
TRINITY_GG_796_c0_g1_i1.p1_FS22A3  ............................................................  109
TRINITY_GG_710_c0_g1_i1.p1_FS38I3  ............................................................  109
TRINITY_GG_734_c0_g1_i1.p1_RC65I3  ............................................................  109
TRINITY_GG_802_c0_g1_i1.p1_RC22A3  ............................................................  109
TRINITY_GG_758_c0_g1_i2.p1_RC75I3  ............................................................  109
```

```
TRINITY_GG_730_c0_g1_i1.p1_FS18A3  ............................................................  109
TRINITY_GG_756_c0_g1_i1.p1_FS30I3  ............................................................  109
TRINITY_GG_730_c0_g1_i2.p1_RC71I3  ............................................................  109

Consensus                          NERELLNSMNISQPTVVFVSKKGLQKILNVQKKLPIIQKIIIMDSKTDYQGFQSMYTFVT  169
TRINITY_GG_752_c0_g1_i1.p1_FS26A3  ............................................................  169
TRINITY_GG_731_c0_g1_i1.p1_RC41A3  ............................................................  169
TRINITY_GG_754_c0_g1_i1.p1_RC104I3 ............................................................  169
TRINITY_GG_807_c0_g1_i1.p1_RC29A3  ............................................................  169
TRINITY_GG_806_c0_g1_i1.p1_RC45A3  ............................................................  169
TRINITY_GG_751_c0_g1_i1.p1_FS35I3  ............................................................  169
TRINITY_GG_741_c0_g1_i1.p1_FS21A3  ............................................................  169
TRINITY_GG_796_c0_g1_i1.p1_FS22A3  ............................................................  169
TRINITY_GG_710_c0_g1_i1.p1_FS38I3  ............................................................  169
TRINITY_GG_734_c0_g1_i1.p1_RC65I3  ............................................................  169
TRINITY_GG_802_c0_g1_i1.p1_RC22A3  ............................................................  169
TRINITY_GG_758_c0_g1_i2.p1_RC75I3  ............................................................  169
TRINITY_GG_730_c0_g1_i1.p1_FS18A3  ............................................................  169
TRINITY_GG_756_c0_g1_i1.p1_FS30I3  ............................................................  169
TRINITY_GG_730_c0_g1_i2.p1_RC71I3  ............................................................  169

Consensus                          SHLPPGFNEYDFVPESFDRDKTIALIMNSSGSTGLPKGVALPHRTACVRFSHARDPIFGN  229
TRINITY_GG_752_c0_g1_i1.p1_FS26A3  ............................................................  229
TRINITY_GG_731_c0_g1_i1.p1_RC41A3  ............................................................  229
TRINITY_GG_754_c0_g1_i1.p1_RC104I3 ............................................................  229
TRINITY_GG_807_c0_g1_i1.p1_RC29A3  ............................................................  229
TRINITY_GG_806_c0_g1_i1.p1_RC45A3  ............................................................  229
TRINITY_GG_751_c0_g1_i1.p1_FS35I3  ............................................................  229
TRINITY_GG_741_c0_g1_i1.p1_FS21A3  ............................................................  229
TRINITY_GG_796_c0_g1_i1.p1_FS22A3  ............................................................  229
TRINITY_GG_710_c0_g1_i1.p1_FS38I3  ............................................................  229
TRINITY_GG_734_c0_g1_i1.p1_RC65I3  ............................................................  229
TRINITY_GG_802_c0_g1_i1.p1_RC22A3  ............................................................  229
TRINITY_GG_758_c0_g1_i2.p1_RC75I3  ............................................................  229
TRINITY_GG_730_c0_g1_i1.p1_FS18A3  ............................................................  229
TRINITY_GG_756_c0_g1_i1.p1_FS30I3  ............................................................  229
TRINITY_GG_730_c0_g1_i2.p1_RC71I3  ............................................................  229

Consensus                          QIIP-DTAILSVVPFHHGFGMFTTLGYLICGFRVVLMYRFEEELFLRSLQDYKIQSALLV  288
TRINITY_GG_752_c0_g1_i1.p1_FS26A3  ....-.......................................................  288
TRINITY_GG_731_c0_g1_i1.p1_RC41A3  ....-.......................................................  288
TRINITY_GG_754_c0_g1_i1.p1_RC104I3 ....-.......................................................  288
TRINITY_GG_807_c0_g1_i1.p1_RC29A3  ....-.......................................................  288
TRINITY_GG_806_c0_g1_i1.p1_RC45A3  ....-.......................................................  288
TRINITY_GG_751_c0_g1_i1.p1_FS35I3  ....-.......................................................  288
TRINITY_GG_741_c0_g1_i1.p1_FS21A3  ....-.......................................................  288
TRINITY_GG_796_c0_g1_i1.p1_FS22A3  ....-.......................................................  288
TRINITY_GG_710_c0_g1_i1.p1_FS38I3  ....-.......................................................  288
TRINITY_GG_734_c0_g1_i1.p1_RC65I3  ....-.......................................................  288
TRINITY_GG_802_c0_g1_i1.p1_RC22A3  ....-.......................................................  288
TRINITY_GG_758_c0_g1_i2.p1_RC75I3  ....-.......................................................  288
TRINITY_GG_730_c0_g1_i1.p1_FS18A3  ....-.......................................................  288
TRINITY_GG_756_c0_g1_i1.p1_FS30I3  ....-.......................................................  288
TRINITY_GG_730_c0_g1_i2.p1_RC71I3  ....-.......................................................  288

Consensus                          PTLFSFFAKSTLIDKYDLSNLHEIASGGAPLSKEVGEAVAKRFHLPGIRQGYGLTETTSA  348
TRINITY_GG_752_c0_g1_i1.p1_FS26A3  ............................................................  348
TRINITY_GG_731_c0_g1_i1.p1_RC41A3  ............................................................  348
TRINITY_GG_754_c0_g1_i1.p1_RC104I3 ............................................................  348
TRINITY_GG_807_c0_g1_i1.p1_RC29A3  ............................................................  348
TRINITY_GG_806_c0_g1_i1.p1_RC45A3  ............................................................  348
TRINITY_GG_751_c0_g1_i1.p1_FS35I3  ............................................................  348
TRINITY_GG_741_c0_g1_i1.p1_FS21A3  ............................................................  348
TRINITY_GG_796_c0_g1_i1.p1_FS22A3  ............................................................  348
TRINITY_GG_710_c0_g1_i1.p1_FS38I3  ............................................................  348
TRINITY_GG_734_c0_g1_i1.p1_RC65I3  ............................................................  348
TRINITY_GG_802_c0_g1_i1.p1_RC22A3  ............................................................  348
TRINITY_GG_758_c0_g1_i2.p1_RC75I3  ............................................................  348
TRINITY_GG_730_c0_g1_i1.p1_FS18A3  ............................................................  348
TRINITY_GG_756_c0_g1_i1.p1_FS30I3  ............................................................  348
TRINITY_GG_730_c0_g1_i2.p1_RC71I3  ............................................................  348

Consensus                          ILITPEGDDKPGAVGKVVPFFEAKVVDLDTGKTLGVNQRGELCVRGPMIMSGYVNNPEAT  408
TRINITY_GG_752_c0_g1_i1.p1_FS26A3  ............................................................  408
TRINITY_GG_731_c0_g1_i1.p1_RC41A3  ............................................................  408
TRINITY_GG_754_c0_g1_i1.p1_RC104I3 ............................................................  408
TRINITY_GG_807_c0_g1_i1.p1_RC29A3  ............................................................  408
TRINITY_GG_806_c0_g1_i1.p1_RC45A3  ............................................................  408
TRINITY_GG_751_c0_g1_i1.p1_FS35I3  ............................................................  408
TRINITY_GG_741_c0_g1_i1.p1_FS21A3  ............................................................  408
TRINITY_GG_796_c0_g1_i1.p1_FS22A3  ............................................................  408
TRINITY_GG_710_c0_g1_i1.p1_FS38I3  ............................................................  408
TRINITY_GG_734_c0_g1_i1.p1_RC65I3  ............................................................  408
TRINITY_GG_802_c0_g1_i1.p1_RC22A3  ............................................................  408
TRINITY_GG_758_c0_g1_i2.p1_RC75I3  ............................................................  408
TRINITY_GG_730_c0_g1_i1.p1_FS18A3  ............................................................  408
TRINITY_GG_756_c0_g1_i1.p1_FS30I3  ............................................................  408
TRINITY_GG_730_c0_g1_i2.p1_RC71I3  ............................................................  408

Consensus                          NALIDKDGWLHSGDIAYWDEDEHFFIVDRLKSLIKYKGYQVAPAELESILLQHPNIFDAG  468
TRINITY_GG_752_c0_g1_i1.p1_FS26A3  ............................................................  468
TRINITY_GG_731_c0_g1_i1.p1_RC41A3  ............................................................  468
TRINITY_GG_754_c0_g1_i1.p1_RC104I3 ............................................................  468
TRINITY_GG_807_c0_g1_i1.p1_RC29A3  ............................................................  468
TRINITY_GG_806_c0_g1_i1.p1_RC45A3  ............................................................  468
TRINITY_GG_751_c0_g1_i1.p1_FS35I3  ............................................................  468
TRINITY_GG_741_c0_g1_i1.p1_FS21A3  ............................................................  468
TRINITY_GG_796_c0_g1_i1.p1_FS22A3  ............................................................  468
TRINITY_GG_710_c0_g1_i1.p1_FS38I3  ............................................................  468
TRINITY_GG_734_c0_g1_i1.p1_RC65I3  ............................................................  468
TRINITY_GG_802_c0_g1_i1.p1_RC22A3  ............................................................  468
TRINITY_GG_758_c0_g1_i2.p1_RC75I3  ............................................................  468
TRINITY_GG_730_c0_g1_i1.p1_FS18A3  ............................................................  468
TRINITY_GG_756_c0_g1_i1.p1_FS30I3  ............................................................  468
TRINITY_GG_730_c0_g1_i2.p1_RC71I3  ............................................................  468

Consensus                          VAGLPDDDAGELPAAVVVLEHGKTMTEKEIVDYVASQVTTAKKLRGGVVFVDEVPKGLTG  528
TRINITY_GG_752_c0_g1_i1.p1_FS26A3  ............................................................  528
TRINITY_GG_731_c0_g1_i1.p1_RC41A3  ............................................................  528
TRINITY_GG_754_c0_g1_i1.p1_RC104I3 ............................................................  528
TRINITY_GG_807_c0_g1_i1.p1_RC29A3  ............................................................  528
TRINITY_GG_806_c0_g1_i1.p1_RC45A3  ............................................................  528
TRINITY_GG_751_c0_g1_i1.p1_FS35I3  ............................................................  528
TRINITY_GG_741_c0_g1_i1.p1_FS21A3  ............................................................  528
TRINITY_GG_796_c0_g1_i1.p1_FS22A3  ............................................................  528
TRINITY_GG_710_c0_g1_i1.p1_FS38I3  ............................................................  528
TRINITY_GG_734_c0_g1_i1.p1_RC65I3  ............................................................  528
TRINITY_GG_802_c0_g1_i1.p1_RC22A3  ............................................................  528
TRINITY_GG_758_c0_g1_i2.p1_RC75I3  ............................................................  528
TRINITY_GG_730_c0_g1_i1.p1_FS18A3  ............................................................  528
TRINITY_GG_756_c0_g1_i1.p1_FS30I3  ............................................................  528
TRINITY_GG_730_c0_g1_i2.p1_RC71I3  ............................................................  528

Consensus                          KLDARKIREILIKAKKGGKSKL                                        550
TRINITY_GG_752_c0_g1_i1.p1_FS26A3  ......................                                        550
TRINITY_GG_731_c0_g1_i1.p1_RC41A3  ......................                                        550
TRINITY_GG_754_c0_g1_i1.p1_RC104I3 ......................                                        550
TRINITY_GG_807_c0_g1_i1.p1_RC29A3  ......................                                        550
TRINITY_GG_806_c0_g1_i1.p1_RC45A3  ......................                                        550
TRINITY_GG_751_c0_g1_i1.p1_FS35I3  ......................                                        550
TRINITY_GG_741_c0_g1_i1.p1_FS21A3  ......................                                        550
TRINITY_GG_796_c0_g1_i1.p1_FS22A3  ......................                                        550
TRINITY_GG_710_c0_g1_i1.p1_FS38I3  ......................                                        550
TRINITY_GG_734_c0_g1_i1.p1_RC65I3  ......................                                        550
TRINITY_GG_802_c0_g1_i1.p1_RC22A3  ......................                                        550
TRINITY_GG_758_c0_g1_i2.p1_RC75I3  ......................                                        550
TRINITY_GG_730_c0_g1_i1.p1_FS18A3  ......................                                        550
TRINITY_GG_756_c0_g1_i1.p1_FS30I3  ......................                                        550
TRINITY_GG_730_c0_g1_i2.p1_RC71I3  ......................                                        550
```

**Figure S3:** Alignment of luciferase sequences derived from light organ transcriptomes. The inactive green light organ replicate FS34I3 was excluded from the alignment, as it omitted from our analysis due to small library size. After trimming and removal of adaptor sequences from mRNA reads, reference-guided transcriptomes were assembled with Trinity (2.10.0). The longest open reading frames were translated with Transdecoder (5.5.0). To identify the luciferase sequence, we used blastp (2.9.0) to obtain the best match (e-value cutoff: 1E-5) between the protein sequences and the published *Photinus pyralis* amino acid sequence (PPYR_00001). All proteins were aligned with MUSCLE in Geneious (2022.2.2) with 100% sequence identity (length: 550 bp), confirming that no differences in the luciferase coding sequence could explain variation in light color.

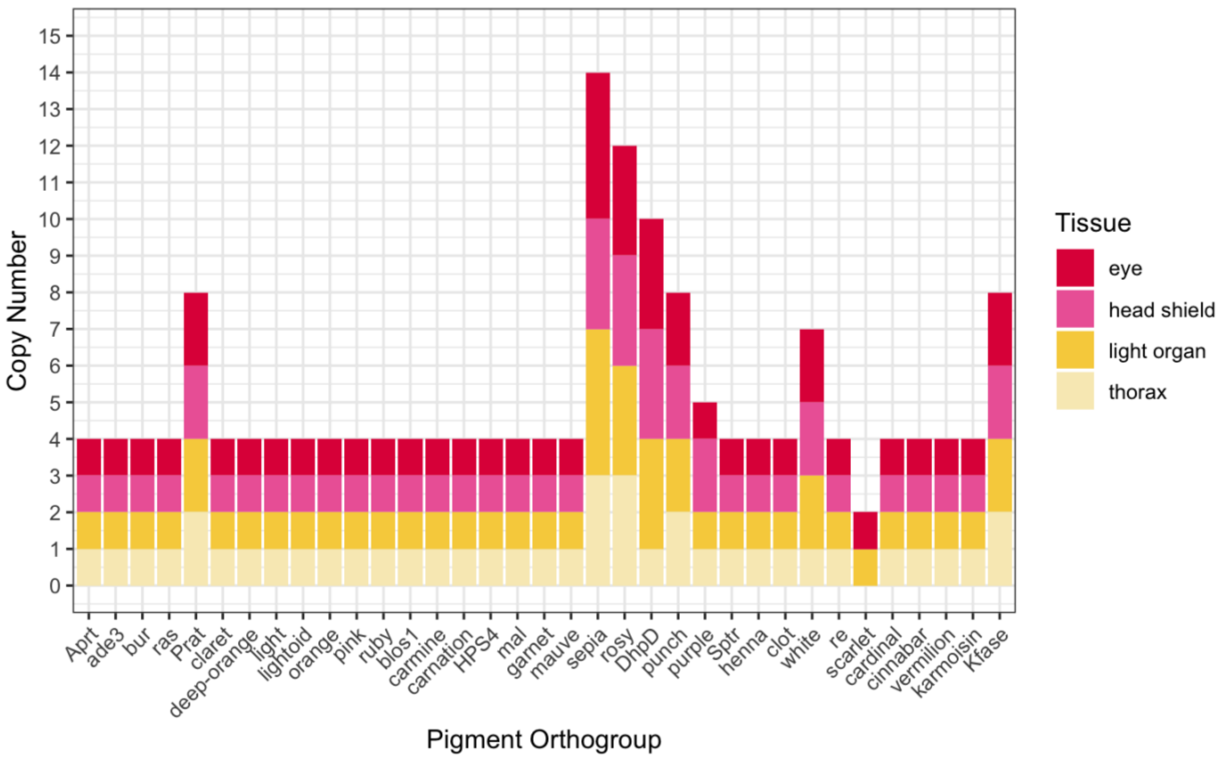


**Figure S5.** Copy number of orthologous pigments sequences across tissues. In *P. pyralis*, pterins appear to have increased copy number compared to other pigment classes.

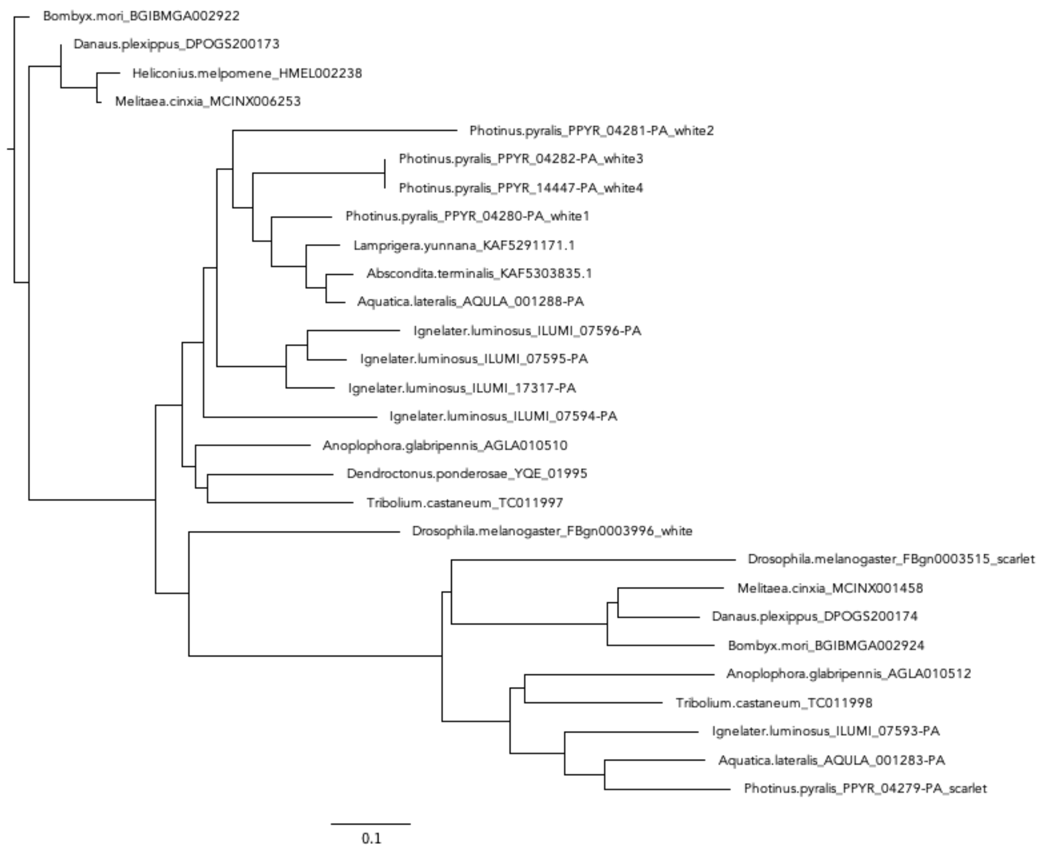


**Figure S6.** Gene tree of white/scarlet gene family. Orthologous protein sequences for each species were aligned with PASTA (v1.8.5) which was trimmed (TrimAl, v1.4.1, default parameters with option -*automated1*) prior to maximum likelihood reconstruction (IQ-TREE, v1.6.12, default parameters with option -m MFP). Gene tree was viewed in FigTree (1.4.4). Nodes represent bootstrap support from 1,000 bootstraps. Tree was rooted with the *Bombyx mori* white ortholog BGIBMGA002922 (Komoto et al., 2009). Clustering patterns between the *Drosophila melanogaster* sequences *scarlet* (FBgn0003515) and *white* (FBgn0003515) with *Photinus pyralis* were used to assign orthology.

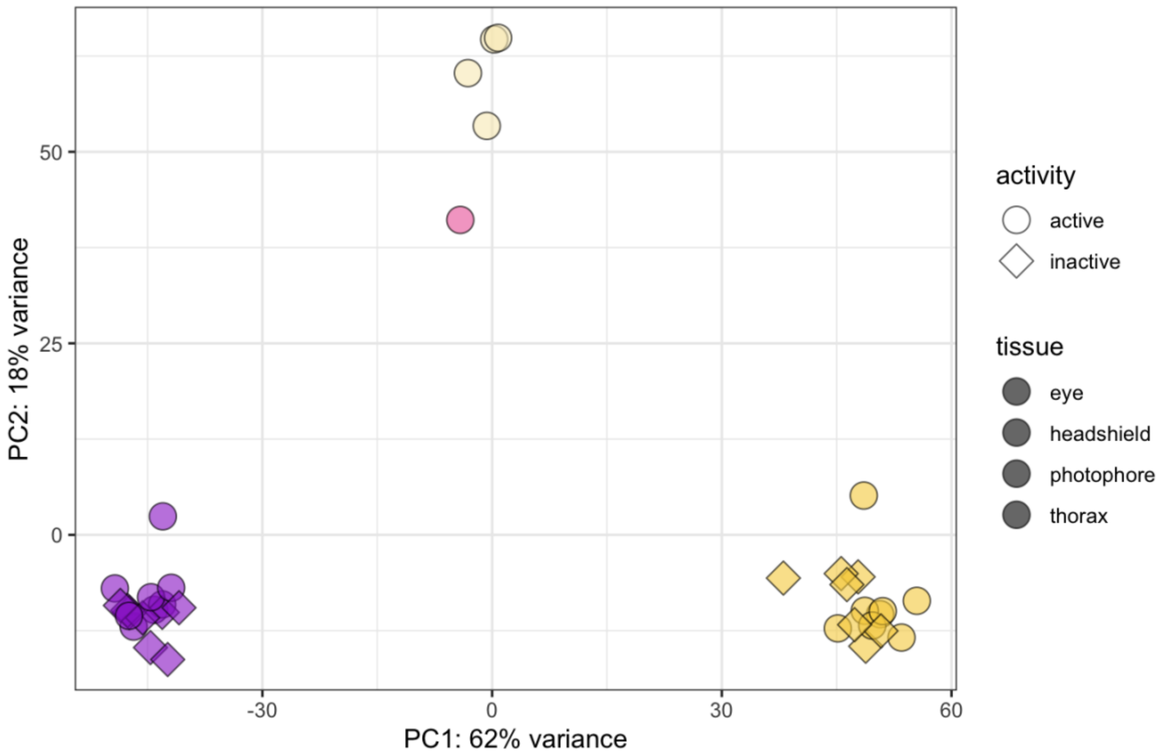


**Figure S7:** PCA of gene expression (total: 12,590 genes) using VST-transformed counts (VST transformation from DESeq2). Tissue type is the major factor driving gene expression differences in *Photinus pyralis*.

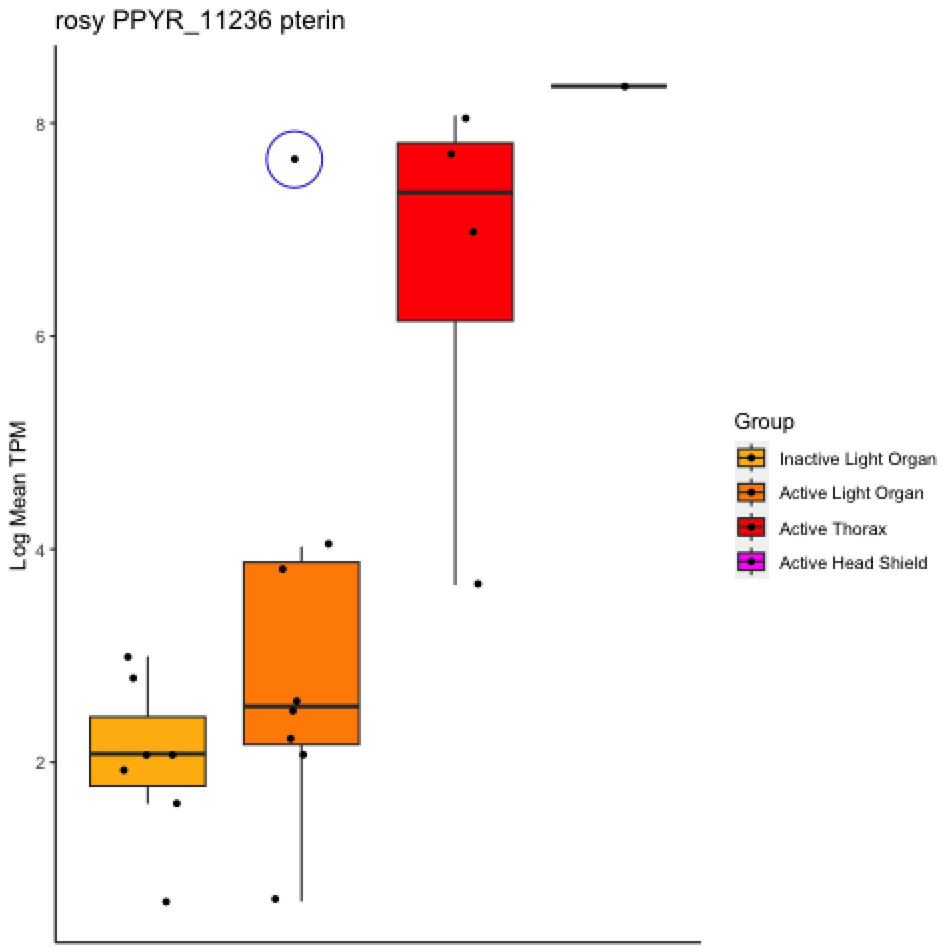


**Figure S8**. Expression of *rosy1* (outlier); differences in expression appear to be driven by one sample (blue circle).

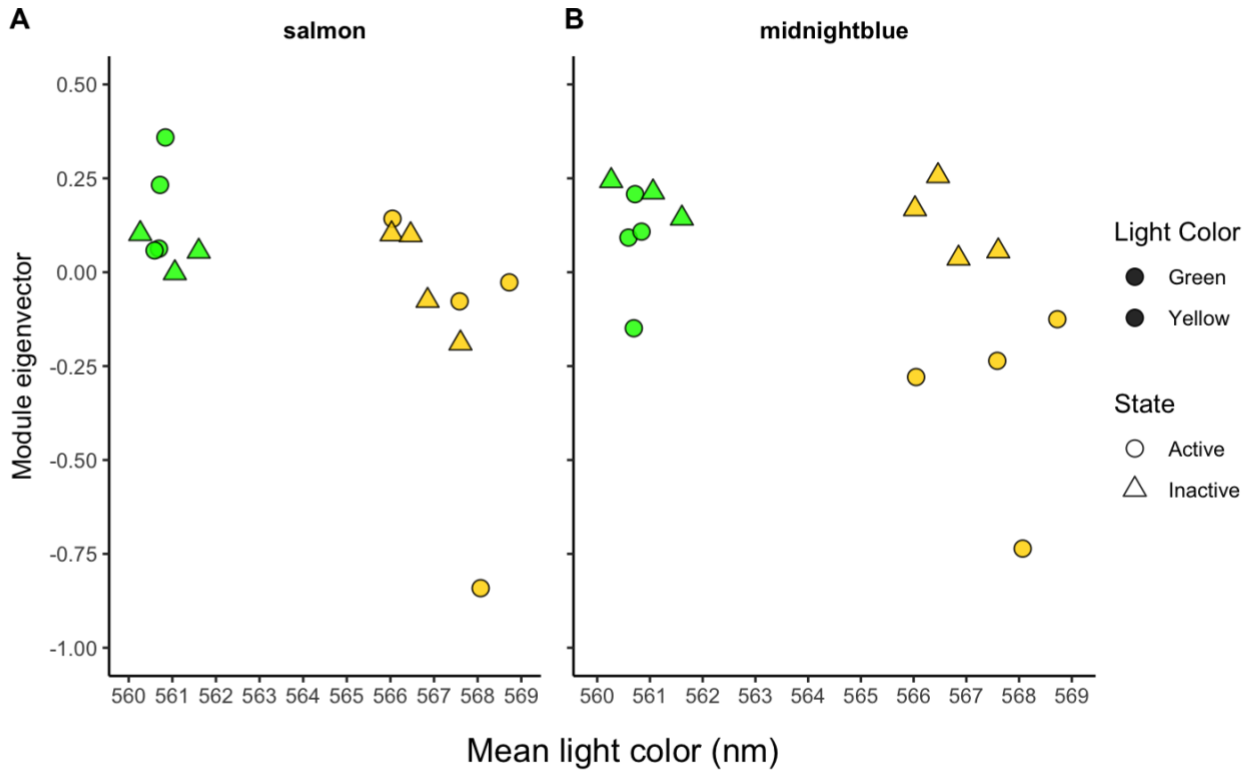


**Figure S9.** Modules associated with light color of *P. pyralis* light organs ($p<0.05$). Shape corresponds to active (circle) inactive (triangle) states. Light color is shown on x-axis as wavelength (nm) while samples are colored according to group (yellow or green). Module eigenvector, or PC1 of module expression, is the y-axis. There is variation within color groups, particularly in the active state of yellow.

**Modules putatively associated with emitted light color**

To identify modules where gene expression was significantly correlated with light color, we used a Pearson correlation between module expression (ME = PC1 or summary value of module genes) and light color (nm) measured from individual fireflies. Using a significance threshold of $p < 0.05$, we detected two modules with marginal significance: M-20 ("midnightblue": p= 0.0508, p-adjusted= 0.8075) and M-30 ("salmon": p=0.0376, p-adjusted=0.8144) (Figure 2.12, Table 2.2). The genes in both modules were more highly expressed in light organs that produced greener light color.

**KEGG Pathway Enrichment Analysis**

To characterize genes differentially expressed between active photic tissues (LO, eyes) with thorax, we performed KEGG enrichment analysis and visualized significant pathways in R with gage (2.52.0) and visualized with Pathview (1.42.0). For DEGs upregulated in either LO or eyes relative to thorax, no pathways survived multiple test correction (Benjamini-Hochberg method) so we considered pathways with $p<0.05$ "of interest." There were nine such pathways upregulated in LO relative to thorax (Table S8), including phagosome (p-value= 0.00580, q-value= 0.47557), lysosome (p-value= 0.02489, q-value= 0.63728), purine metabolism (p-value= 0.01115, q-value= 0.47557), and neuroactive ligand receptor (p-value= 0.02045, q-value= 0.63728) (shown below).

Enrichment of the lysosome pathway could have a role related to pigments, as pigments are lysosomal-related organelles (LROs). We observed the upregulation of genes that encode lysosomal acid hydrolases, as well as membrane proteins and acidification regulators. This was particularly interesting, because intracellular conditions (e.g., pH, redox state, presence of metals and/or proteins) can alter the molecular structure of pigments, resulting in changes to their absorption (Figon et al., 2021). We also observed the upregulation of genes encoding proteins associated with lysosomal trafficking, which contribute to the biogenesis of pigment granules. Notably, this included the AP-3 complex, whose subunits are encoded by members of the granule group *garnet* (Simpson et al., 1997), *ruby* (Kretzschmar et al., 2000; Mullins et al., 2000), *carmine* (Mullins et al., 1999), and *orange* (Mullins et al., 2000).

Purines are involved in a multitude of roles, including biosynthesis of pterins. The purine degradation pathway gives rise to pterins through incorporation guanine (GTP) into the pigment granule; however, guanine can also be directly converted to xanthine, which results in the final breakdown product, uric acid (Vogels & Van der Drift, 1976). Fireflies have abundant levels of uric acid in their LO, which forms the reflective layer to amplify their light signals outward through the clear cuticle (Goh et al., 2013). It remains unclear whether either of these pathways are indicative of pigments, rather suggests that existing pathways required for their synthesis are prominent in LO.

We additionally observed sulfur metabolism (p-value= 0.04221, q-value= 0.67537) and biosynthesis of unsaturated fatty acids (p-value= 0.03987, q-value= 0.03987), which are consistent with luciferin synthesis (Fallon et al., 2016) and luciferase activity (Oba et al., 2003), respectively, as luciferase evolved via gene duplication of fatty acyl-CoA synthase (Fallon et al., 2018).

Among genes upregulated in eyes relative to thorax, we observed the enrichment of phototransduction (p-value= 0.02658, q-value= 1.0000), as expected due to the vision, in addition to Neuroactive ligand-receptor interaction (p-value= 0.00156, q-value= 0.19992). This pathway was also enriched in active LO, suggesting some similarity in physiological response while signaling.

<u>KEGG pathways “of interest” (p-value<0.05) upregulated in LO (LO vs. thorax) (for complete list, see Table S8).</u>

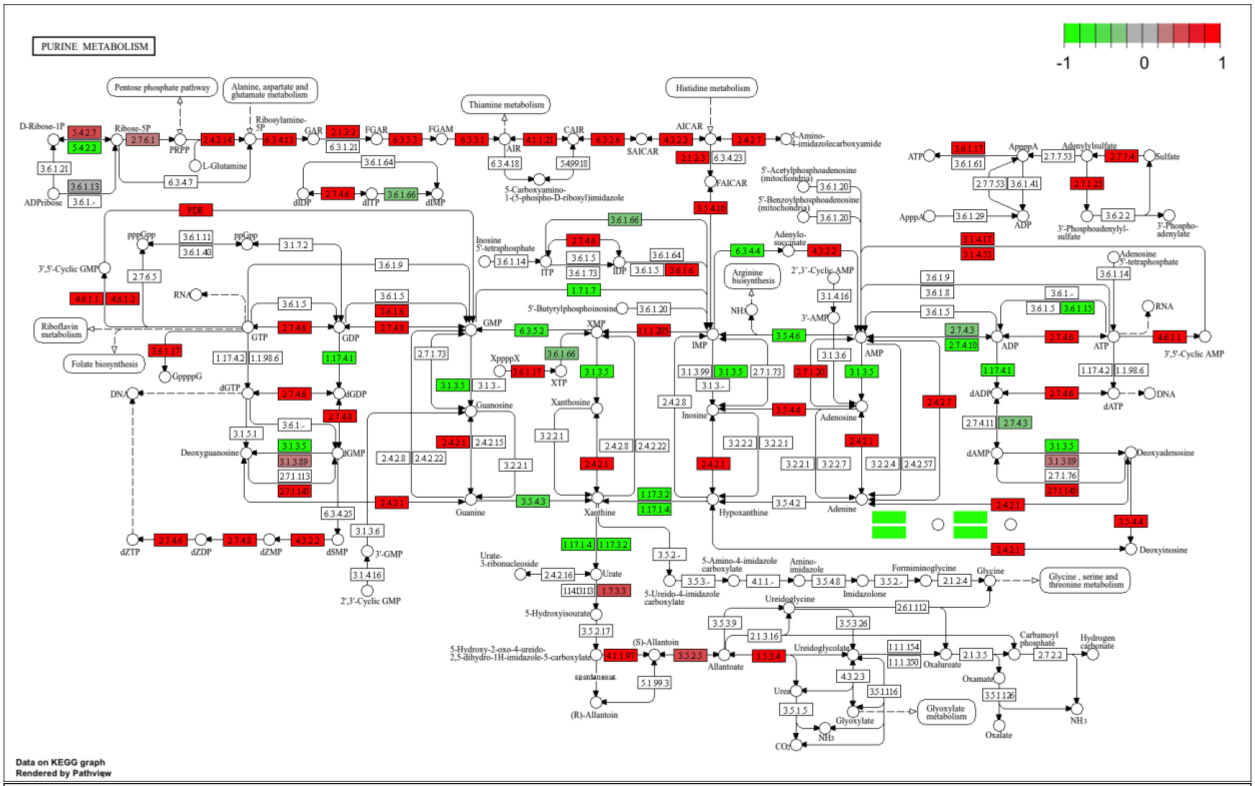

PURINE METABOLISM
-1
0
1
Pentose phosphate pathway
Alanine, aspartate and glutamate metabolism
Thiamine metabolism
Histidine metabolism
D-Ribose-1P
Ribose-5P
PRPP
L-Glutamine
Ribosylamine-5P
GAR
FGAR
FGAM
AIR
CAIR
SAICAR
AICAR
FAICAR
ADPRibose
dIDP
dITP
dIMP
ppGpp
pppGpp
3',5'-Cyclic GMP
RNA
Riboflavin metabolism
Folate biosynthesis
GppppG
GTP
GDP
GMP
XMP
IMP
dGTP
dGDP
DNA
Deoxyguanosine
dGMP
Guanosine
Xanthosine
Inosine
Adenosine
Guanine
Xanthine
Hypoxanthine
Adenine
dZTP
dZDP
dZMP
dSMP
3'-GMP
2',3'-Cyclic GMP
XTP
ITP
IDP
Arginine biosynthesis
AMP
ADP
ATP
dADP
dATP
dAMP
Deoxyadenosine
Deoxyinosine
3'-AMP
2',3'-Cyclic AMP
3',5'-Cyclic AMP
Adenylo-succinate
AppppA
ApppA
Adenylylsulfate
Sulfate
3'-Phosphoadenylyl-sulfate
3'-Phospho-adenylate
Urate
Allantoate
Urea
Glyoxylate
Glyoxylate metabolism
Oxalureate
Oxamate
Oxalate
Carbamoyl phosphate
Hydrogen carbonate
Glycine
Glycine, serine and threonine metabolism
Imidazolone
Ureidoglycine
Ureidoglycolate
(S)-Allantoin
(R)-Allantoin
5-Hydroxyisourate
spontaneous
CO2
NH3
Data on KEGG graph
Rendered by Pathview


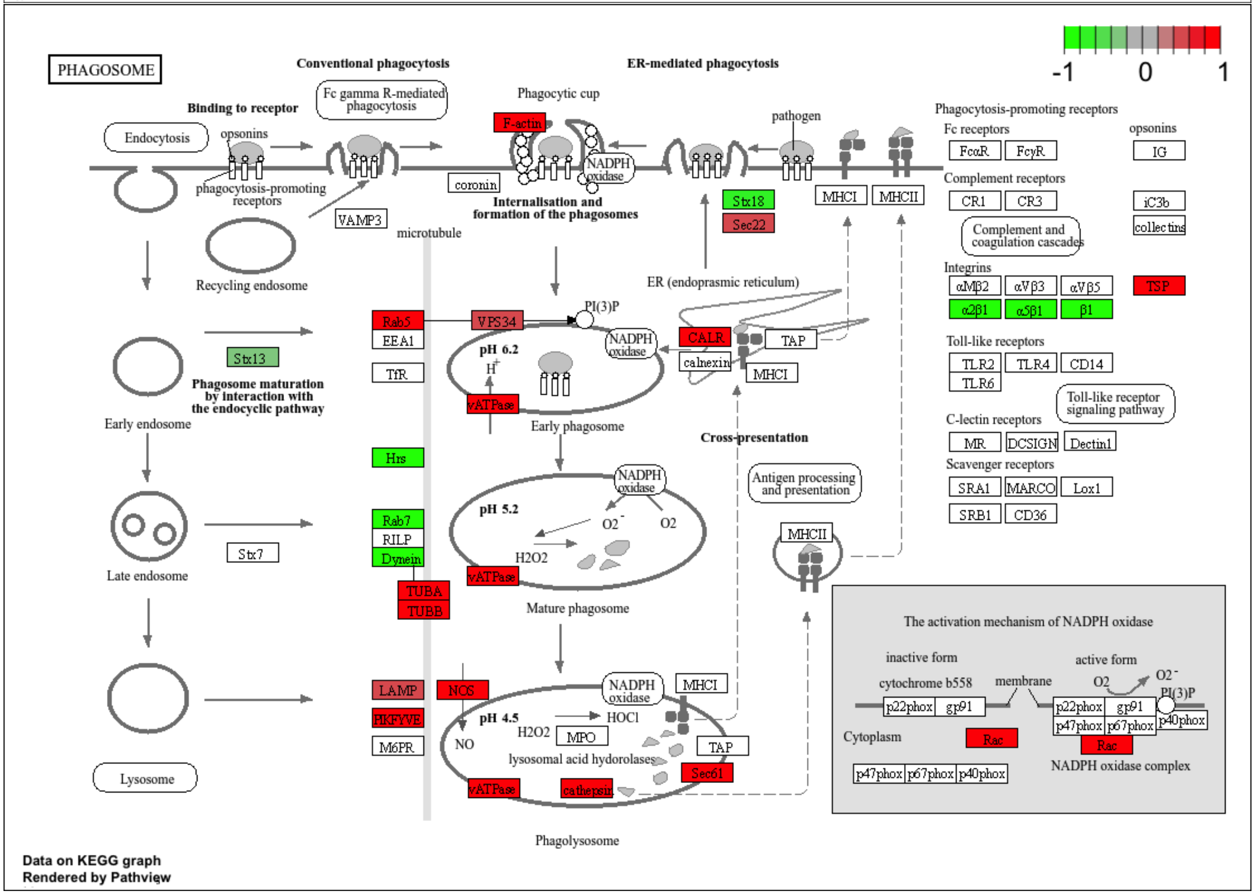

PHAGOSOME
-1
0
1
Conventional phagocytosis
ER-mediated phagocytosis
Binding to receptor
Fc gamma R-mediated phagocytosis
Phagocytic cup
pathogen
Endocytosis
opsonins
phagocytosis-promoting receptors
F-actin
NADPH oxidase
coronin
VAMP3
Internalisation and formation of the phagosomes
Stx18
Sec22
MHCI
MHCII
microtubule
Recycling endosome
ER (endoplasmic reticulum)
Rab5
EEA1
TfR
VPS34
PI(3)P
pH 6.2
CALR
calnexin
TAP
Stx13
Phagosome maturation by interaction with the endocyclic pathway
Early endosome
vATPase
Early phagosome
Cross-presentation
Hrs
Antigen processing and presentation
Rab7
RILP
Dynein
Stx7
Late endosome
pH 5.2
O2
H2O2
TUBA
TUBB
Mature phagosome
LAMP
NOS
PIKFYVE
M6PR
pH 4.5
HOCl
MPO
NO
lysosomal acid hydrolases
cathepsin
Sec61
Lysosome
Phagolysosome
Phagocytosis-promoting receptors
Fc receptors
FcαR
FcγR
IG
Complement receptors
CR1
CR3
iC3b
collectins
Complement and coagulation cascades
Integrins
αMβ2
αVβ3
αVβ5
TSP
α2β1
α5β1
β1
Toll-like receptors
TLR2
TLR4
CD14
TLR6
Toll-like receptor signaling pathway
C-lectin receptors
MR
DCSIGN
Dectin1
Scavenger receptors
SRA1
MARCO
Lox1
SRB1
CD36
The activation mechanism of NADPH oxidase
inactive form
active form
cytochrome b558
membrane
p22phox
gp91
p47phox
p67phox
p40phox
Cytoplasm
Rac
NADPH oxidase complex
Data on KEGG graph
Rendered by Pathview

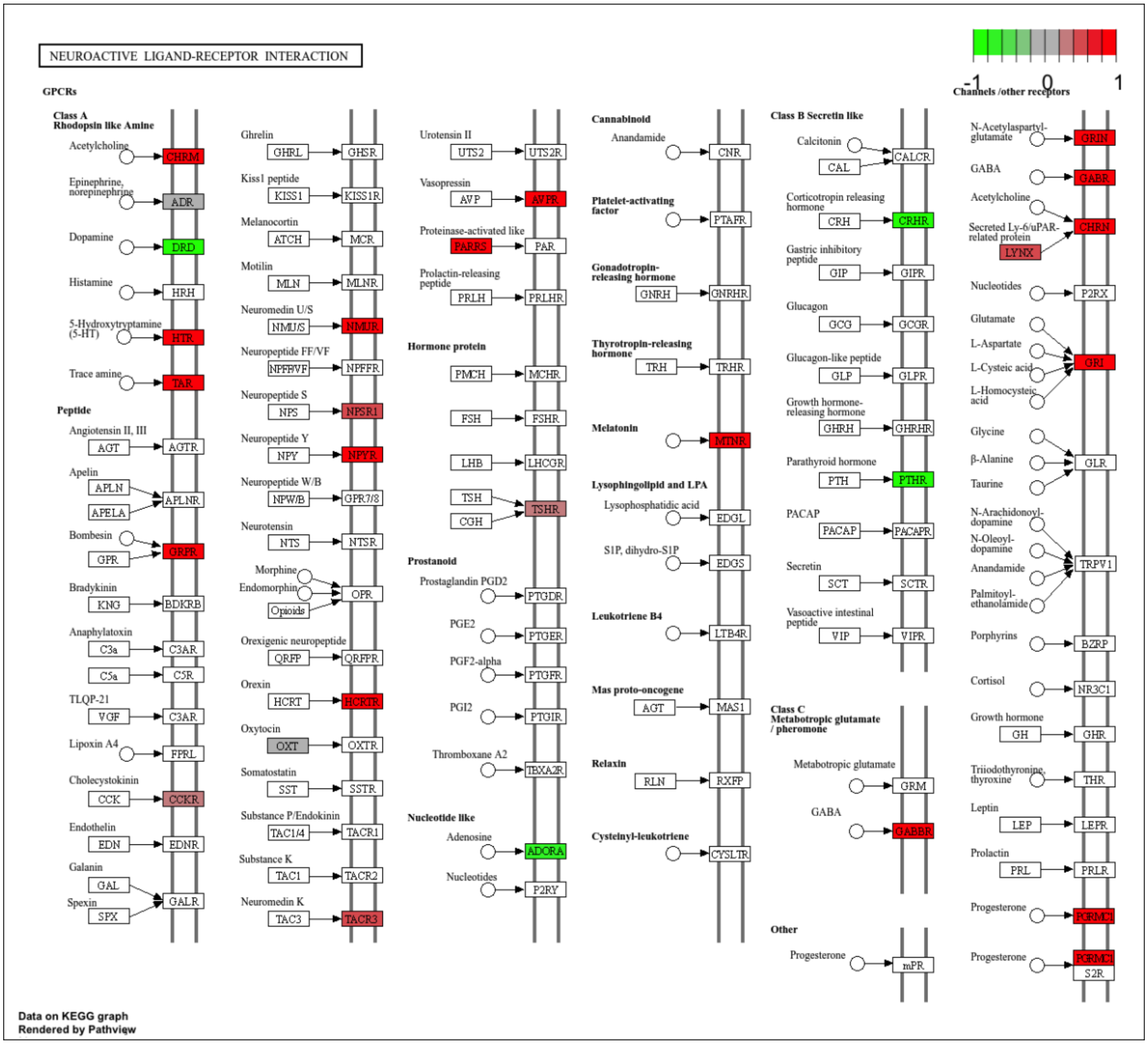
NEUROACTIVE LIGAND-RECEPTOR INTERACTION
-1
0
1
GPCRs
Class A
Rhodopsin like Amine
Acetylcholine
CHRM
Epinephrine, norepinephrine
ADR
Dopamine
DRD
Histamine
HRH
5-Hydroxytryptamine (5-HT)
HTR
Trace amine
TAR
Peptide
Angiotensin II, III
AGT
AGTR
Apelin
APLN
APELA
APLNR
Bombesin
GPR
GRPR
Bradykinin
KNG
BDKRB
Anaphylatoxin
C3a
C3AR
C5a
C5R
TLQP-21
VGF
C3AR
Lipoxin A4
FPRL
Cholecystokinin
CCK
CCKR
Endothelin
EDN
EDNR
Galanin
GAL
Spexin
SPX
GALR
Ghrelin
GHRL
GHSR
Kiss1 peptide
KISS1
KISS1R
Melanocortin
ATCH
MCR
Motilin
MLN
MLNR
Neuromedin U/S
NMU/S
NMUR
Neuropeptide FF/VF
NPFF/VF
NPFFR
Neuropeptide S
NPS
NPSR1
Neuropeptide Y
NPY
NPYR
Neuropeptide W/B
NPW/B
GPR7/8
Neurotensin
NTS
NTSR
Morphine
Endomorphin
Opioids
OPR
Orexigenic neuropeptide
QRFP
QRFPR
Orexin
HCRT
HCRTR
Oxytocin
OXT
OXTR
Somatostatin
SST
SSTR
Substance P/Endokinin
TAC1/4
TACR1
Substance K
TAC1
TACR2
Neuromedin K
TAC3
TACR3
Urotensin II
UTS2
UTS2R
Vasopressin
AVP
AVPR
Proteinase-activated like
PARRS
PAR
Prolactin-releasing peptide
PRLH
PRLHR
Hormone protein
PMCH
MCHR
FSH
FSHR
LHB
LHCGR
TSH
CGH
TSHR
Prostanoid
Prostaglandin PGD2
PTGDR
PGE2
PTGER
PGF2-alpha
PTGFR
PGI2
PTGIR
Thromboxane A2
TBXA2R
Nucleotide like
Adenosine
ADORA
Nucleotides
P2RY
Cannabinoid
Anandamide
CNR
Platelet-activating factor
PTAFR
Gonadotropin-releasing hormone
GNRH
GNRHR
Thyrotropin-releasing hormone
TRH
TRHR
Melatonin
MTNR
Lysophingolipid and LPA
Lysophosphatidic acid
EDGL
S1P, dihydro-S1P
EDGS
Leukotriene B4
LTB4R
Mas proto-oncogene
AGT
MAS1
Relaxin
RLN
RXFP
Cysteinyl-leukotriene
CYSLTR
Class B Secretin like
Calcitonin
CAL
CALCR
Corticotropin releasing hormone
CRH
CRHR
Gastric inhibitory peptide
GIP
GIPR
Glucagon
GCG
GCGR
Glucagon-like peptide
GLP
GLPR
Growth hormone-releasing hormone
GHRH
GHRHR
Parathyroid hormone
PTH
PTHR
PACAP
PACAP
PACAPR
Secretin
SCT
SCTR
Vasoactive intestinal peptide
VIP
VIPR
Class C
Metabotropic glutamate / pheromone
Metabotropic glutamate
GRM
GABA
GABBR
Other
Progesterone
mPR
Channels /other receptors
N-Acetylaspartyl-glutamate
GRIN
GABA
GABR
Acetylcholine
Secreted Ly-6/uPAR-related protein
LYNX
CHRN
Nucleotides
P2RX
Glutamate
L-Aspartate
L-Cysteic acid
L-Homocysteic acid
GRI
Glycine
β-Alanine
Taurine
GLR
N-Arachidonoyl-dopamine
N-Oleoyl-dopamine
Anandamide
Palmitoyl-ethanolamide
TRPV1
Porphyrins
BZRP
Cortisol
NR3C1
Growth hormone
GH
GHR
Triiodothyronine, thyroxine
THR
Leptin
LEP
LEPR
Prolactin
PRL
PRLR
Progesterone
PGRMC1
Progesterone
PGRMC1
S2R
Data on KEGG graph
Rendered by Pathview

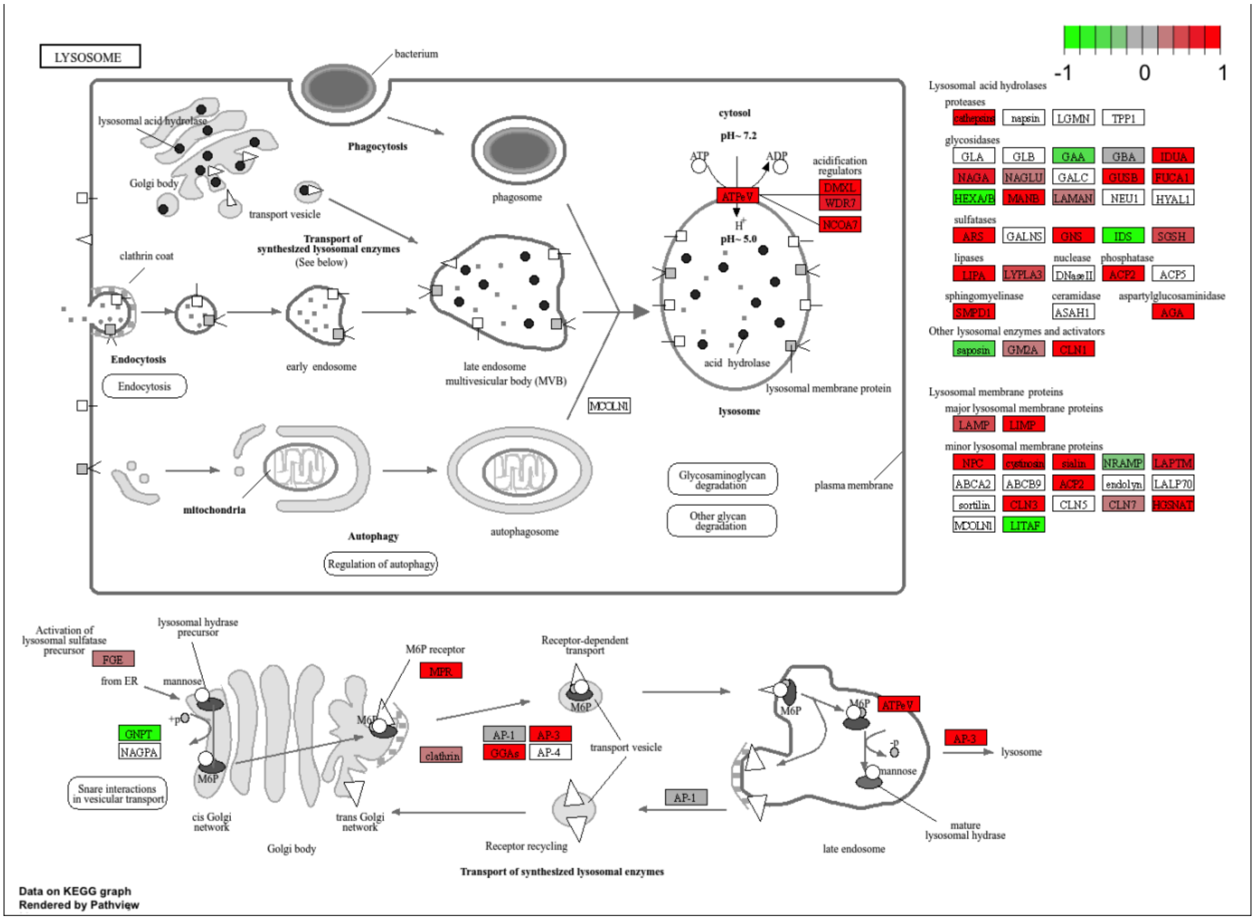
LYSOSOME
bacterium
-1
0
1
lysosomal acid hydrolase
Golgi body
transport vesicle
Phagocytosis
phagosome
cytosol
pH~ 7.2
ATP
ADP
acidification regulators
ATPeV
DMXL
WDR7
NCOA7
pH~ 5.0
Transport of synthesized lysosomal enzymes
(See below)
clathrin coat
Endocytosis
early endosome
late endosome
multivesicular body (MVB)
MCOLN1
acid hydrolase
lysosomal membrane protein
lysosome
mitochondria
Autophagy
autophagosome
Regulation of autophagy
Glycosaminoglycan degradation
Other glycan degradation
plasma membrane
Lysosomal acid hydrolases
proteases
cathepsins
napsin
LGMN
TPP1
glycosidases
GLA
GLB
GAA
GBA
IDUA
NAGA
NAGLU
GALC
GUSB
FUCA1
HEXA/B
MANB
LAMAN
NEU1
HYAL1
sulfatases
ARS
GALNS
GNS
IDS
SGSH
lipases
nuclease
phosphatase
LIPA
LYPLA3
DNaseII
ACP2
ACP5
sphingomyelinase
ceramidase
aspartylglucosaminidase
SMPD1
ASAH1
AGA
Other lysosomal enzymes and activators
saposin
GM2A
CLN1
Lysosomal membrane proteins
major lysosomal membrane proteins
LAMP
LIMP
minor lysosomal membrane proteins
NPC
cystinosin
sialin
NRAMP
LAPTM
ABCA2
ABCB9
ACP2
endolyn
LALP70
sortilin
CLN3
CLN5
CLN7
HGSNAT
MCOLN1
LITAF
Activation of lysosomal sulfatase precursor
FGE
lysosomal hydrase precursor
from ER
mannose
GNPT
NAGPA
Snare interactions in vesicular transport
M6P
cis Golgi network
Golgi body
M6P receptor
MPR
clathrin
trans Golgi network
AP-1
AP-3
GGAs
AP-4
Receptor-dependent transport
transport vesicle
Receptor recycling
AP-1
late endosome
ATPeV
AP-3
lysosome
mannose
mature lysosomal hydrase
Transport of synthesized lysosomal enzymes
Data on KEGG graph
Rendered by Pathview

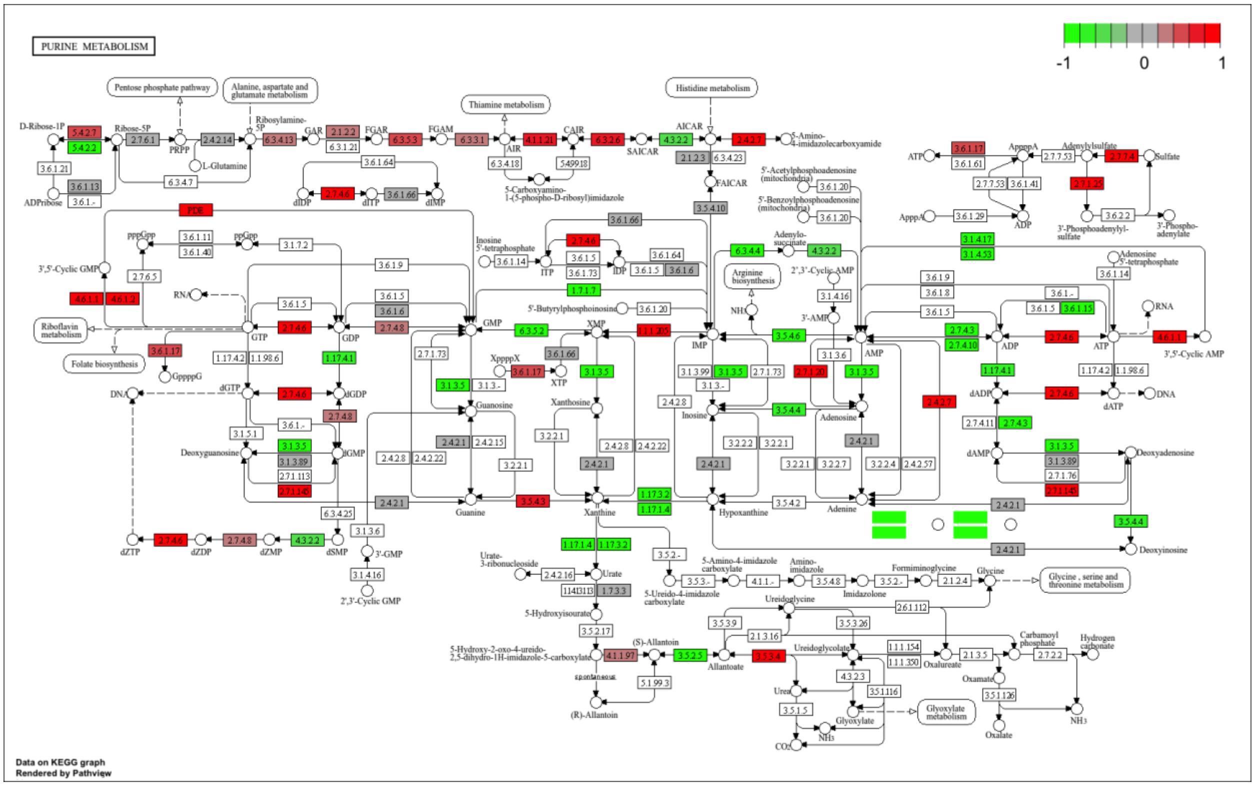


KEGG pathways “of interest” (p-value<0.05) upregulated in eyes (eyes vs. thorax)

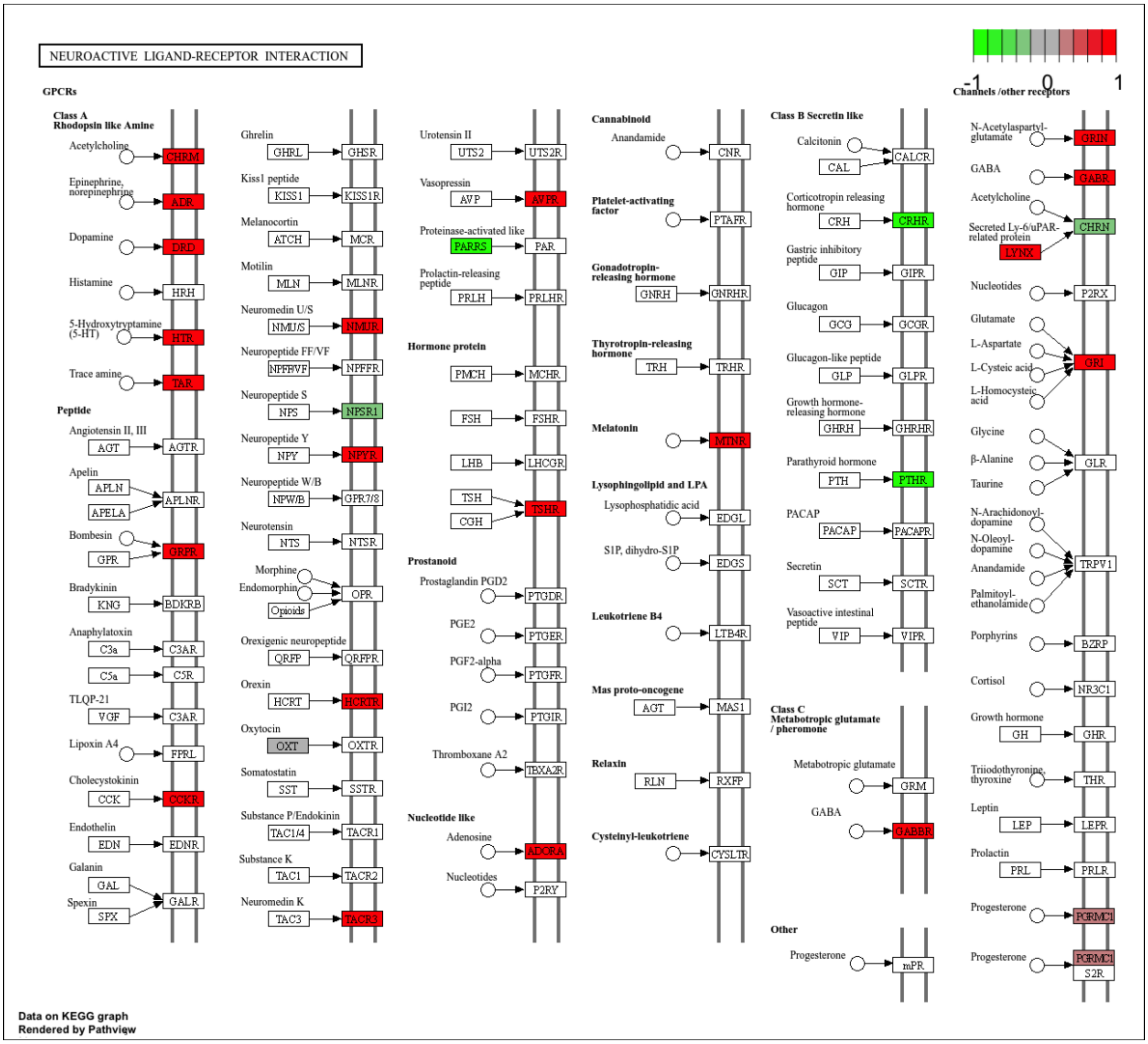
NEUROACTIVE LIGAND-RECEPTOR INTERACTION
-1
0
1
GPCRs
Class A
Rhodopsin like Amine
Acetylcholine
CHRM
Epinephrine, norepinephrine
ADR
Dopamine
DRD
Histamine
HRH
5-Hydroxytryptamine (5-HT)
HTR
Trace amine
TAR
Peptide
Angiotensin II, III
AGT
AGTR
Apelin
APLN
APELA
APLNR
Bombesin
GPR
GRPR
Bradykinin
KNG
BDKRB
Anaphylatoxin
C3a
C3AR
C5a
C5R
TLQP-21
VGF
C3AR
Lipoxin A4
FPRL
Cholecystokinin
CCK
CCKR
Endothelin
EDN
EDNR
Galanin
GAL
Spexin
SPX
GALR
Ghrelin
GHRL
GHSR
Kiss1 peptide
KISS1
KISS1R
Melanocortin
ATCH
MCR
Motilin
MLN
MLNR
Neuromedin U/S
NMU/S
NMUR
Neuropeptide FF/VF
NPFF/VF
NPFFR
Neuropeptide S
NPS
NPSR1
Neuropeptide Y
NPY
NPYR
Neuropeptide W/B
NPW/B
GPR7/8
Neurotensin
NTS
NTSR
Morphine
Endomorphin
Opioids
OPR
Orexigenic neuropeptide
QRFP
QRFPR
Orexin
HCRT
HCRTR
Oxytocin
OXT
OXTR
Somatostatin
SST
SSTR
Substance P/Endokinin
TAC1/4
TACR1
Substance K
TAC1
TACR2
Neuromedin K
TAC3
TACR3
Urotensin II
UTS2
UTS2R
Vasopressin
AVP
AVPR
Proteinase-activated like
PARRS
PAR
Prolactin-releasing peptide
PRLH
PRLHR
Hormone protein
PMCH
MCHR
FSH
FSHR
LHB
LHCGR
TSH
CGH
TSHR
Prostanoid
Prostaglandin PGD2
PTGDR
PGE2
PTGER
PGF2-alpha
PTGFR
PGI2
PTGIR
Thromboxane A2
TBXA2R
Nucleotide like
Adenosine
ADORA
Nucleotides
P2RY
Cannabinoid
Anandamide
CNR
Platelet-activating factor
PTAFR
Gonadotropin-releasing hormone
GNRH
GNRHR
Thyrotropin-releasing hormone
TRH
TRHR
Melatonin
MTNR
Lysophingolipid and LPA
Lysophosphatidic acid
EDGL
S1P, dihydro-S1P
EDGS
Leukotriene B4
LTB4R
Mas proto-oncogene
AGT
MAS1
Relaxin
RLN
RXFP
Cysteinyl-leukotriene
CYSLTR
Class B Secretin like
Calcitonin
CAL
CALCR
Corticotropin releasing hormone
CRH
CRHR
Gastric inhibitory peptide
GIP
GIPR
Glucagon
GCG
GCGR
Glucagon-like peptide
GLP
GLPR
Growth hormone-releasing hormone
GHRH
GHRHR
Parathyroid hormone
PTH
PTHR
PACAP
PACAP
PACAPR
Secretin
SCT
SCTR
Vasoactive intestinal peptide
VIP
VIPR
Class C
Metabotropic glutamate / pheromone
Metabotropic glutamate
GRM
GABA
GABBR
Other
Progesterone
mPR
Channels /other receptors
N-Acetylaspartyl-glutamate
GRIN
GABA
GABR
Acetylcholine
Secreted Ly-6/uPAR-related protein
LYNX
CHRN
Nucleotides
P2RX
Glutamate
L-Aspartate
L-Cysteic acid
L-Homocysteic acid
GRI
Glycine
β-Alanine
Taurine
GLR
N-Arachidonoyl-dopamine
N-Oleoyl-dopamine
Anandamide
Palmitoyl-ethanolamide
TRPV1
Porphyrins
BZRP
Cortisol
NR3C1
Growth hormone
GH
GHR
Triiodothyronine, thyroxine
THR
Leptin
LEP
LEPR
Prolactin
PRL
PRLR
Progesterone
PGRMC1
Progesterone
PGRMC1
S2R
Data on KEGG graph
Rendered by Pathview

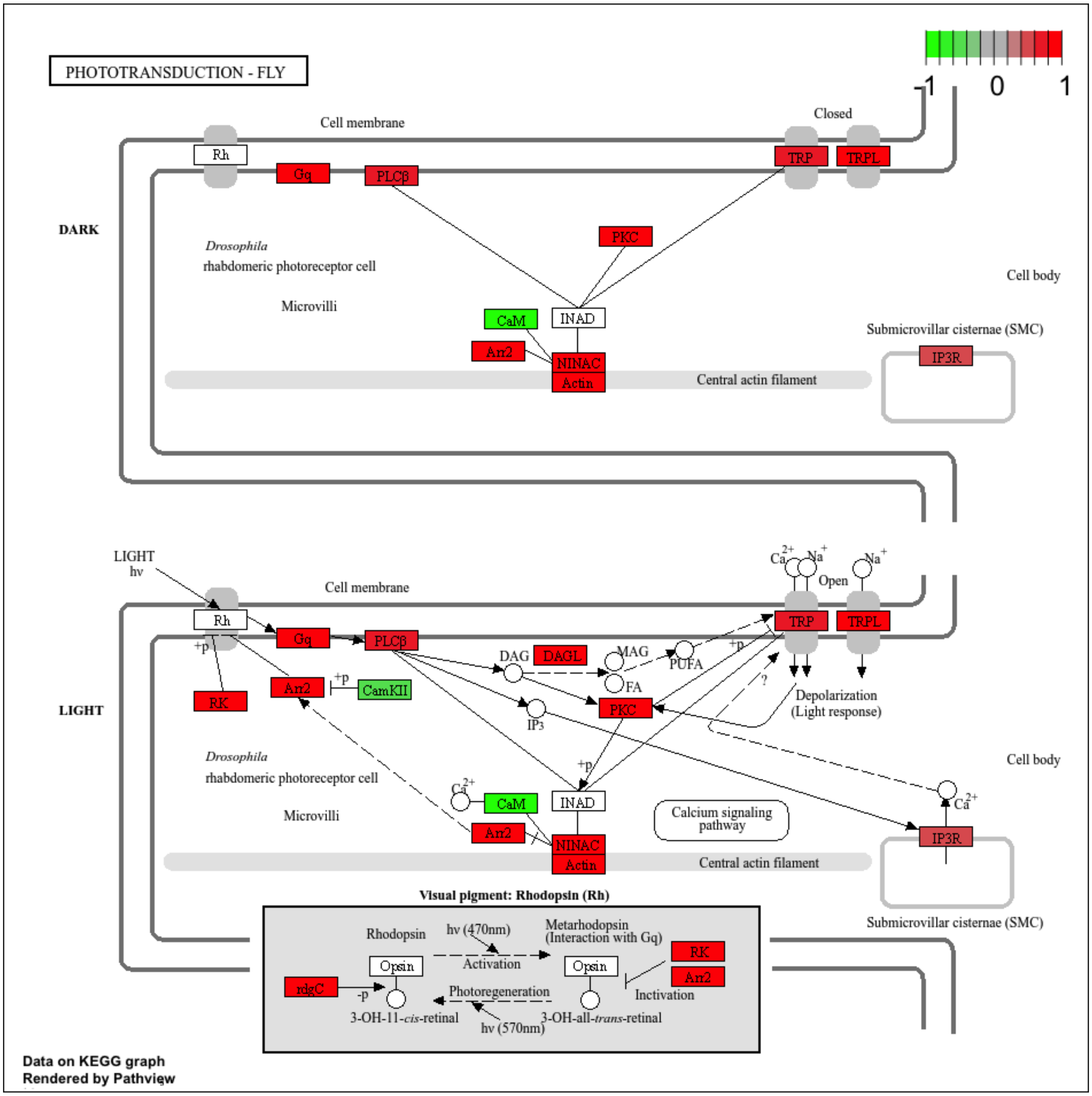


### Network analysis: Module 27 "Red"

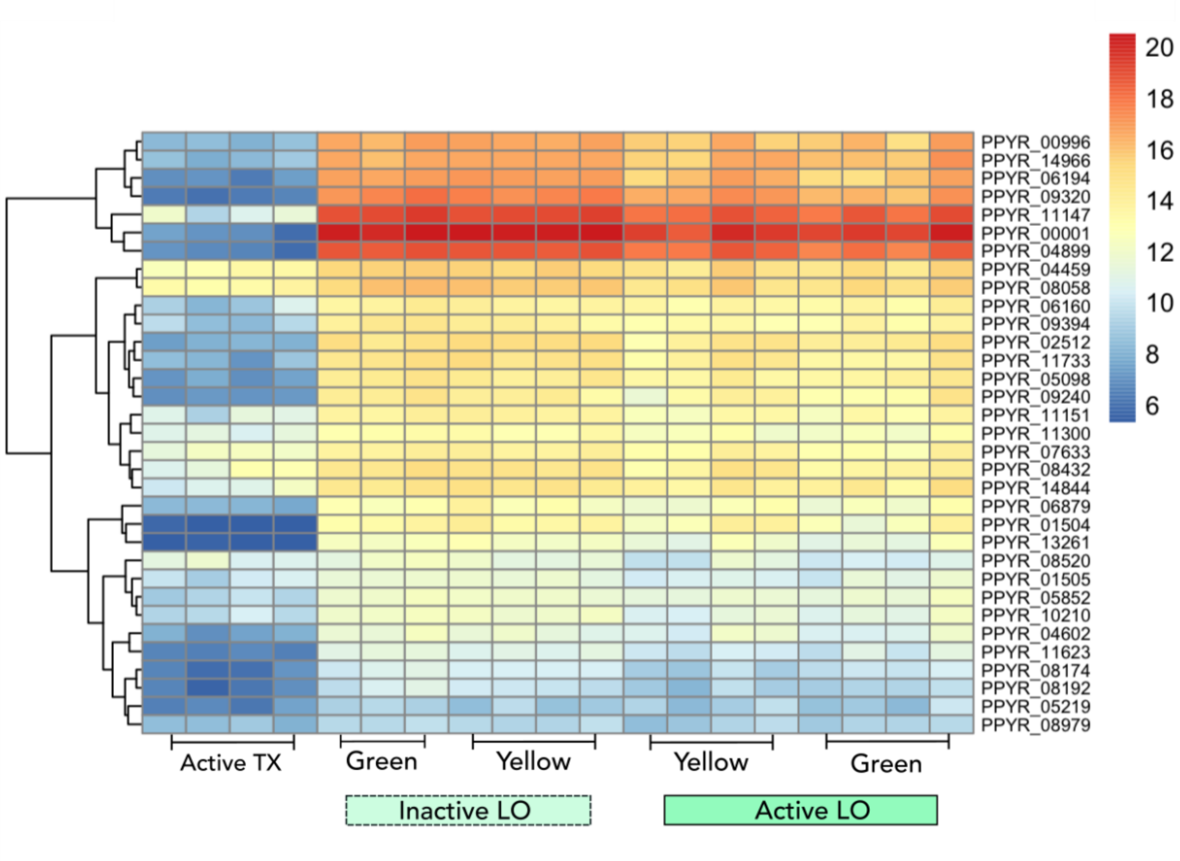


The 546 genes in module "red" (M-27) were significantly ($p<0.05$) associated with inactive light organs, including one pigment gene: sepia5 (pterin). *Sepia5* was expressed in all tissues, with highest expression in the eye (mean TPM±SD: x̄=781.25±208.47, N=16), followed by light organ (x̄= 351.67±101.43, N=15) and thorax (x̄=196.25±98.21, N=4). Notably, 34 bioluminescence genes (of 73 in total) were present in this module (Figure S11), including luciferase (PPYR_00001), which we identified as a hub (Table S18). Indeed, Module "Red" was significantly enriched for LO genes from Fallon et al. (2018) (FDR=1.95E-25), in addition to "photic" (FDR= 5.04E-10) and LO-specific (FDR=1.90E-22) DEG (Contrast 2). Also present in this module were genes involved in phototransduction, including: *chaoptin* (PPYR_00272) and *retinol-binding protein pinta-like* (PPYR_01227) as well as those regulating circadian rhythm: *cryptochrome-1-like* (PPYR_08961), *potassium voltage-gated channel protein Shaker* (PPYR_14129), and six copies of *takeout-like* (Table S17). This was consistent with our GO analysis (Table S20), which found circadian rhythm (p-value=0.00120, FDR=1.00000), in addition to terms related to acidity: vacuolar acidification (p-value=0.00024, FDR=0.37144), pH reduction (p-value=0.00024, 1.00000), and response to alkaline pH (p-value=0.00182, FDR=1.00000) in addition to purine ribonucleoside monophosphate metabolism (p-value=0.00136, FDR=1.00000) and purine biosynthesis (p-value=0.00264, FDR=1.00000).
